# The formation of K_V_2.1 macro-clusters is required for sex-specific differences in L-type Ca_V_1.2 clustering and function in arterial myocytes

**DOI:** 10.1101/2023.06.27.546725

**Authors:** Collin Matsumoto, Samantha C. O’Dwyer, Declan Manning, Gonzalo Hernandez-Hernandez, Paula Rhana, Zhihui Fong, Daisuke Sato, Colleen E. Clancy, Nicholas C. Vierra, James S. Trimmer, L. Fernando Santana

## Abstract

In arterial myocytes, the canonical function of voltage-gated Ca_V_1.2 and K_V_2.1 channels is to induce myocyte contraction and relaxation through their responses to membrane depolarization, respectively. Paradoxically, K_V_2.1 also plays a sex-specific role by promoting the clustering and activity of Ca_V_1.2 channels. However, the impact of K_V_2.1 protein organization on Ca_V_1.2 function remains poorly understood. We discovered that K_V_2.1 forms micro-clusters, which can transform into large macro-clusters when a critical clustering site (S590) in the channel is phosphorylated in arterial myocytes. Notably, female myocytes exhibit greater phosphorylation of S590, and macro-cluster formation compared to males. Contrary to current models, the activity of K_V_2.1 channels seems unrelated to density or macro-clustering in arterial myocytes. Disrupting the K_V_2.1 clustering site (K_V_2.1_S590A_) eliminated K_V_2.1 macro-clustering and sex-specific differences in Ca_V_1.2 cluster size and activity. We propose that the degree of K_V_2.1 clustering tunes Ca_V_1.2 channel function in a sex-specific manner in arterial myocytes.

## Introduction

Activation of dihydropyridine-sensitive, voltage-gated L-type Ca_V_1.2 channels plays a crucial role in the development of myogenic tone (*1*), a process of autoregulation that enables arteries to regulate their diameter in response to changes in intravascular pressure (*2*). This mechanism, independent of neural or hormonal influences, is critical to maintain constant blood flow despite changes in blood pressure.

The current model of the myogenic response proposes that mechanical stretch of the membrane leads to the activation of TRPC6 (*3*), TRPM4 (*4*), and TRPP1 (PKD2) (*5*) channels, which depolarizes arterial myocytes, activating Ca_V_1.2 channels (*1*). Activation of a single or a small cluster of Ca_V_1.2 channels results in a local increase in intracellular Ca^2+^ concentration ([Ca^2+^]_i_) termed a “Ca_V_1.2 sparklet” (*6-8*). Summation of multiple Ca_V_1.2 sparklets leads to a global increase in [Ca^2+^]_i_ that triggers muscle contraction.

Ca_V_1.2 channels form clusters in the plasma membrane via a stochastic self-assembly mechanism (*9*). Ca_V_1.2 channels within these clusters can gate cooperatively in response to Ca^2+^-driven physical interactions of adjacent channels (*10, 11*). Ca_V_1.2 channels in this configuration allow for larger Ca^2+^ influx compared to random, independent openings of individual channels. In vascular smooth muscle, cooperative gating of Ca_V_1.2 channels has been estimated to contribute up to ∼50% of Ca^2+^ influx during the development of myogenic tone (*12*).

One route of negative feedback regulation of membrane depolarization and Ca^2+^ influx via Ca_V_1.2 channels occurs through the depolarization-mediated activation of delayed rectifier voltage-gated K_V_2.1 channels (*13, 14*). In its canonical role, K_V_2.1 proteins in arterial smooth muscle cells form ion conducting voltage-gated K^+^ channels whose activation results in membrane potential hyperpolarization, thereby affecting myocyte [Ca^2+^]_i_ and myogenic tone (*13*). Until recently, the accepted role of K_V_2.1 protein in arterial myocytes was to form K^+^ conducting channels. However, our recent work suggests that only about 1% of the K_V_2.1 channels in the plasma membrane of arterial smooth muscle are conductive (*15*). Indeed, a growing body of evidence suggests that K_V_2.1 proteins have dual conductive and structural roles in the surface membrane of smooth muscle cells and neurons (*15-17*).

In neurons and HEK293T cells, K_V_2.1 is expressed in large macro-clusters (*16, 18-23*). A 26 amino acid region within the C-terminus of the channel called the proximal restriction and clustering (PRC) domain was determined to be responsible for this expression pattern (*24*). The high-density clustering of K_V_2.1 channels is influenced by phosphorylation of serine residues within the PRC domain (*25-27*). Additionally, in heterologous expression systems, the majority of K_V_2.1 channels within macro-clusters are considered non-conductive (*18, 19, 28*). Little to no channel activity was detectable within K_V_2.1 clusters, whereas currents could be recorded in areas with diffuse K_V_2.1 expression (*19*). One of the structural roles of K_V_2.1 is to promote clustering of Ca_V_1.2 channels, thus increasing the probability of Ca_V_1.2-to-Ca_V_1.2 interactions within these clusters (*16, 17*).

Both, the conductive and structural roles of K_V_2.1, depend on the level of expression of this protein, which in arterial myocytes varies with sex (*15*). In female myocytes, where expression of K_V_2.1 protein is higher than in male myocytes, K_V_2.1 has both conductive and structural roles. Female myocytes have larger Ca_V_1.2 clusters, [Ca^2+^]_i_, and myogenic tone than male myocytes. In contrast, in male myocytes, K_V_2.1 channels regulate membrane potential, but not Ca_V_1.2 channel clustering.

Based on these data, a model was proposed in which K_V_2.1 function varies with sex (*15*). In males, K_V_2.1 channels primarily control membrane potential, but in female myocytes K_V_2.1 plays dual electrical and Ca_V_1.2 clustering roles. Currently, it is unclear whether the conductive and structural functions of K_V_2.1 in native arterial myocytes rely on its clustering ability, and if this relationship is sex-dependent.

In this study, we tested the hypothesis that conductive and structural roles of K_V_2.1 channels in male and female arterial myocytes depend on their capacity to form clusters in studies of wild-type (WT) and S586A mutant rat K_V_2.1 channels expressed in heterologous cells, and in arterial myocytes from a novel gene edited knock-in mouse expressing the S590A mutation. We focused on serine 586 within the PRC domain (amino acids 573-598) of rat K_V_2.1 because a point mutation changing this amino acid to a non-phosphorylatable alanine decreases the K_V_2.1 clustering phenotype (*24*). This corresponds to a S590A point mutation in the mouse K_V_2.1 channel. Our data show that K_V_2.1 is expressed into large macro-clusters composed of micro-clusters that can only be resolved with super-resolution microscopy. The K_V_2.1_S586A_ point mutation nearly eliminated K_V_2.1 macro-clusters but had a minimal impact on micro-clusters. Notably, we find that K_V_2.1 channel function is not dependent on its ability to form macro-clusters in arterial myocytes of either sex. Rather, K_V_2.1 macro-clustering enhances Ca_V_1.2 channel clusters and activity. Based on these results, we propose a new model in which macro-clustering of K_V_2.1 in arterial myocytes alters Ca_V_1.2 channel organization and function in a sex-specific manner but has no impact on its conductive function.

## Results

### K_V_2.1 macro-clusters are composed of micro-clusters of K_V_2.1 and are declustered by the K_V_2.1_S586A_ mutation

We began our study by determining the spatial distribution in heterologous cells of exogenously expressed wild-type rat K_V_2.1 (K_V_2.1_WT_) channels and K_V_2.1 channels in which serine at position 586 was mutated to an alanine (K_V_2.1_S586A_) using confocal and super-resolution ground state depletion (GSD) microscopy (**Figure 1**). Both channels were tagged at their N-terminus with the red-shifted fluorescent protein mScarlet.

**Figure 1.**
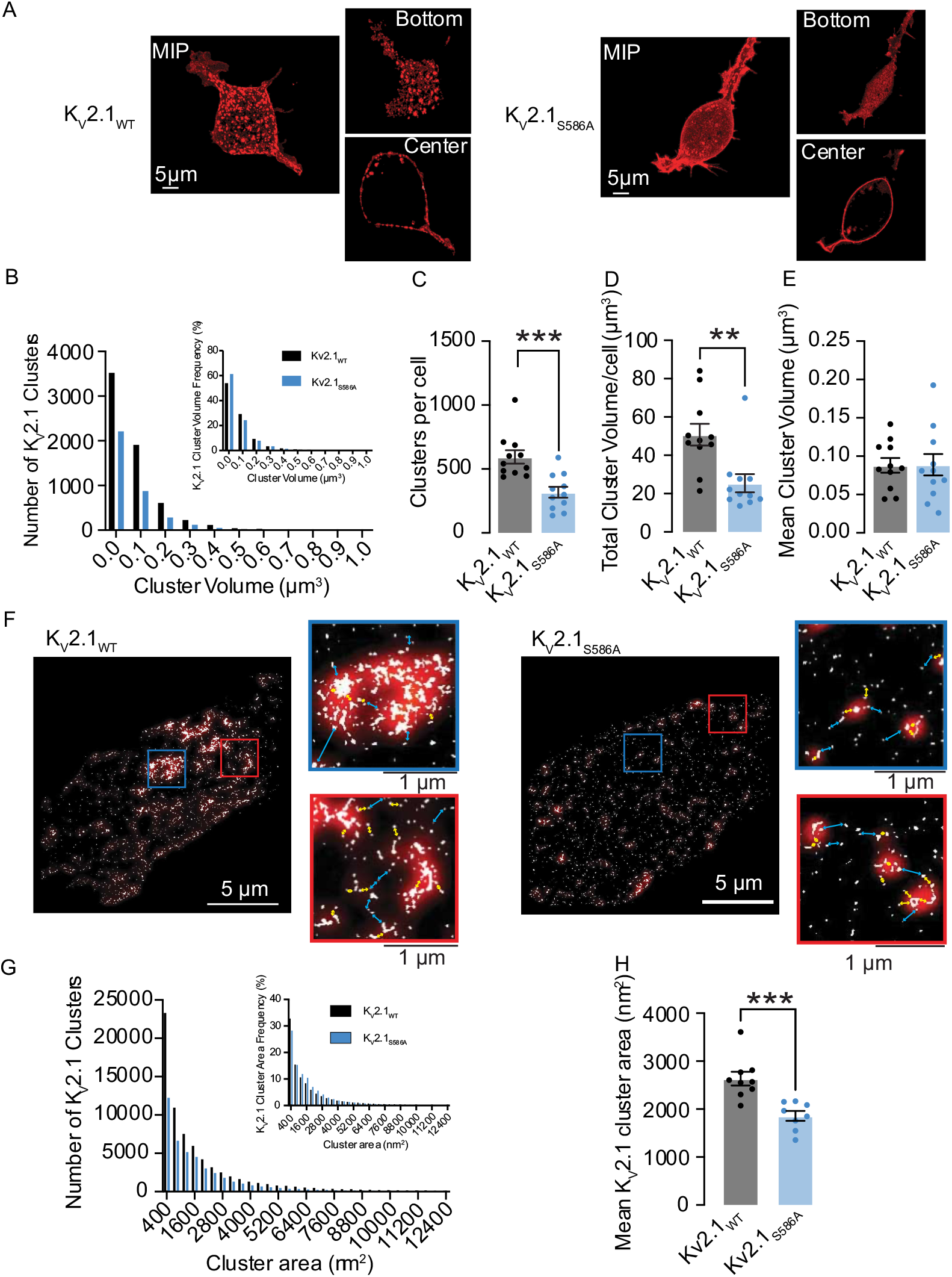
K_V_2.1 macro-clusters are declustered into micro-clusters with the K_V_2.1_S586A_ point mutation. (**A**) Confocal maximum intensity projection images of HEK293T cells transfected with K_V_2.1_WT_ (left) or K_V_2.1_S586A_ (right) tagged with mScarlet. Insets show single plane images of the bottom or center of each cell. (**B**) Number of K_V_2.1 cluster volumes of K_V_2.1_WT_ (black) and K_V_2.1_S586A_ (blue) in transfected HEK293T cells. Insets quantify relative frequency histograms as a percentage of K_V_2.1 volumes. Plots representing (**C**) number of clusters per cell, (**D**) total cluster volume per cell, and (**E**) mean cluster volumes. (**F**) Representative super-resolution GSD images of immunolabeled K_V_2.1 channels in transfected HEK293T cells. Insets show 4 μm^2^ regions of interest. Red overlay depicts Gaussian blur. Cyan lines indicate distances of greater than 200 nm while yellow arrows represent distances less than 200 nm. (**G**) Number of clusters of K_V_2.1_WT_ (black) and K_V_2.1_S586A_ (blue) by cluster area in transfected HEK293T cells. Insets quantify relative frequency histograms as a percentage of K_V_2.1 areas. (**H**) Summary plot of mean K_V_2.1 cluster areas from super-resolution images. *P < 0.05, **P < 0.01, ***P < 0.001. Error bars indicate mean ± SEM.

**Figure 1A** shows confocal maximum intensity projection images of 3D reconstructions of representative HEK293T cells expressing K_V_2.1_WT_ (left) and K_V_2.1_S586A_ (right). To the right of each image, we show single plane images from the bottom (top panel) and center (bottom panel) of each cell. **Figure 1B-E** shows a quantitative analysis of the number and volume of K_V_2.1 clusters from these 3D confocal images. The frequency distributions of K_V_2.1_WT_ (black) and K_V_2.1_S586A_ (blue) cluster volumes could be fit with a single exponential function. Of note, the number of clusters were smaller in most volume bins for the K_V_2.1_S586A_ mutation compared to K_V_2.1_WT_. For example, in the same number of cells (n = 7), we detected a total of 6,344 K_V_2.1_WT_ clusters, but only 3,335 K_V_2.1_S586A_ clusters. The number of clusters per cell was 594.4 ± 52.7 in K_V_2.1_WT_ (median = 542 clusters) and 318.2 ± 42.7 clusters in K_V_2.1_S586A_ (median = 299 clusters) (P = 0.0003) (**Figure 1C**). The total cluster volume per cell of K_V_2.1_WT_ was 50.8 ± 5.7 μm^3^ (median = 47.8 μm^3^) compared to 25.4 ± 4.7 μm^3^ in cells expressing K_V_2.1_S586A_ (median = 19.8 μm^3^) (P = 0.001) (**Figure 1D**). Interestingly, the mean cluster volumes were not significantly different between K_V_2.1_WT_ channels at 0.09 ± 0.01 μm^3^ (median = 0.08 μm^3^) and K_V_2.1_S586A_ at 0.09 ± 0.01 μm^3^ (median = 0.08 μm^3^) (**Figure 1E**). These data suggest that K_V_2.1_WT_ channels are expressed into clusters and that K_V_2.1_S586A_ expression is more diffuse and exhibits a more uniform distribution.

**Figure 1F** provides representative super-resolution ground-state depletion (GSD) TIRF images (lateral resolution ≈ 40 nm) from representative cells expressing K_V_2.1_WT_ or K_V_2.1_S586A_. Note that data are provided on an area basis since we are imaging a single plane footprint of the cell in contrast to capturing multiple Z-slices. We also provide two regions of interest (boxed areas) by each image. K_V_2.1_WT_ and K_V_2.1_S586A_ channels are organized into clusters of varied sizes (**Figure 1G**), and consistent with the lower resolution confocal data, the distribution of Kv2.1_S586A_ channel clusters was more diffuse than that of K_V_2.1_WT_. It should be noted that the regions of interest (ROI) of our GSD images reveal that the macro-clusters observed at the confocal level are, in fact, made up of numerous micro-clusters.

To increase our confidence in this observation, we utilized a Gaussian blur (shown in red) to decrease the resolution of the GSD signal to match that of confocal microscopy. Cyan arrows indicate distances greater than 200 nm between signals produced by GSD, while yellow arrows indicate distances less than 200 nm (**Figure 1F, insets**). Notably, our GSD image with a Gaussian blur accurately reproduced the clustering phenotype of K_V_2.1_WT_ at the confocal level, providing further evidence that macro-clusters are composed of micro-clusters. Although we do not observe the large macro-clusters in K_V_2.1_S586A_ expressing cells, we can still resolve K_V_2.1 micro-clusters.

The frequency distribution of the cluster areas of K_V_2.1_WT_ and K_V_2.1_S586A_ obtained from GSD imaging could both be fit with an exponential function (**Figure 1G**). The mean cluster area of K_V_2.1_WT_ was 2634 ± 143 nm^2^ (median = 2568 nm^2^), larger than that of K_V_2.1_S586A_ channels which was 1860 ± 104 nm^2^ (median = 1907 nm^2^) (P = 0.0003) (**Figure 1H**). This is likely due to the absence of larger clusters in cells expressing K_V_2.1_S586A_. Notably, cluster density was 41 ± 5 clusters per micron and 29 ± 4 clusters per micron for K_V_2.1 and K_V_2.1_S586A_, respectively.

Our finding that the size distributions of K_V_2.1_WT_ and K_V_2.1_S586A_ clusters could be described by exponential functions, the hallmark of a *Poisson* process, suggests that these clusters are formed stochastically (*29*). To test this hypothesis, we implemented the stochastic modeling approach employed by Sato et al. (*9*) to determine whether we could reproduce our cluster distributions and make testable predictions regarding plasma membrane protein organization. As shown in **Supplemental Figure 1**, our stochastic self-assembly model effectively reproduced the steady-state size distributions that we measured for K_V_2.1_WT_ and K_V_2.1_S586A_ proteins embedded in the surface membrane of HEK293T cells. The parameters used in the model are summarized in **Supplemental 1C**. These *in silico* data suggest that in HEK293T cells, K_V_2.1_WT_ has a higher probability of nucleation (i.e., P_n_) and cluster growth (i.e., P_g_) than K_V_2.1_S586A_ channels. The probability of channel removal (P_R_) was similar.

Analysis of our confocal microscopy data showed that in HEK293T cells, K_V_2.1_WT_ is expressed in clusters of different sizes but do in fact form large macro-clusters. Using this confocal microscopy analysis, we sought to define the size of a K_V_2.1 macro-cluster. We began with the mean cluster volume of K_V_2.1_WT_ generated in **Figure 1E**. Our analysis provided a mean cluster volume of 0.09 μm^3^. Assuming the volume of a cluster is spherical, we extrapolated the diameter of the macro-cluster to be 560 nm. The standard deviation of these measurements was 0.32. Thus, the mean minus 2 standard deviations provides a lower end limit with a 95% confidence and aligned with the lateral resolution of our confocal microscopy. Accordingly, we define the lower limit of a macro-cluster as a cluster that is larger than 0.03 μm^3^ or 326 nm in diameter, with clusters smaller than this classified as micro-clusters.

### K_V_2.1_S590A_ mutation does not affect overall K_V_2.1 channel expression but declusters smooth muscle K_V_2.1 channels in a sex-specific manner

Next, we investigated whether, as in heterologous expression system (i.e., **Figure 1**), K_V_2.1 channels cluster in arterial myocytes and whether this clustering could be disrupted via mutation of critical amino acids in the PRC domain. To do this, we used Crispr/CAS gene editing to generate a knock-in mouse line expressing the S590A point mutation, corresponding to the S586A mutation in rat K_V_2.1, at the KCNB1 locus of a C57/BL6J mouse (see Methods section for a full description of how these mice were generated).

We isolated, fixed, permeabilized, and labeled arterial myocytes from both male and female K_V_2.1_WT_ and K_V_2.1_S590A_ knock-in mice. We double labeled with wheat germ agglutinin-488 (WGA488) to identify the sarcolemma and K_V_2.1-specific antibodies in male and female K_V_2.1_WT_ and K_V_2.1_S590A_ myocytes using confocal microscopy. **Figure 2A and 2G** show representative 3D images of fixed mesenteric smooth muscle cells labeled with WGA488 and immunolabeled for K_V_2.1.

**Figure 2.**
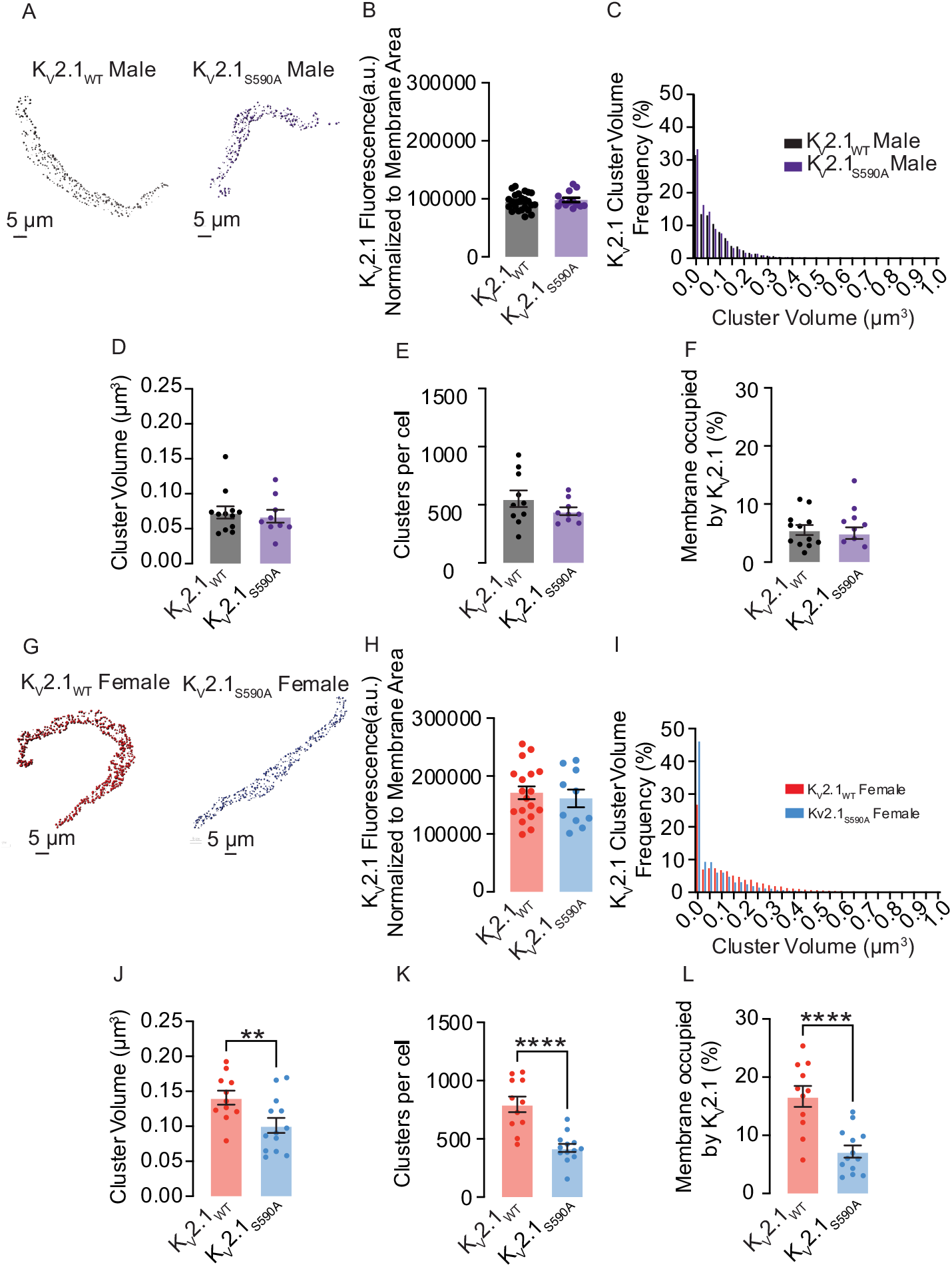
The K_V_2.1_S590A_ mutation declusters K_V_2.1 channels in arterial smooth muscle in a sex-specific manner. (**A, G**) Representative maximum projection images of K_V_2.1 channel clusters at the surface membrane of K_V_2.1_WT_ male (**A**, left), K_V_2.1_S590A_ male (**A**, right), K_V_2.1_WT_ female (**G**, left) and K_V_2.1_S590A_ female (**G**, right) myocytes. Quantification of immunofluorescence normalized to cell membrane area of labeled K_V_2.1 in male (**B**) and female **(H**) myocytes. (**C, I**) Relative frequency as a percentage of K_V_2.1 cluster volumes from K_V_2.1_WT_ male (**C**, black), K_V_2.1_S590A_ male (**C**, purple), K_V_2.1_WT_ female (**I**, red) and K_V_2.1_S590A_ female (**I**, blue) myocytes. Summary data of K_V_2.1 clusters in myocytes showing (**D, J**) mean cluster volumes, (**E, K**) clusters per cell, and (**F, L**) percent of the surface membrane occupied by K_V_2.1 channels in male and female myocytes. *P<0.05, **P<0.01, ***P<0.001, ****P<0.0001. Error bars indicate mean ± SEM.

We investigated whether the K_V_2.1_S590A_ mutation leads to altered expression levels of the channel in arterial myocytes. Our analysis showed that total, cell-wide K_V_2.1-associated fluorescence normalized to cell membrane area was similar in sex-matched K_V_2.1_WT_ and K_V_2.1_S590A_ myocytes (**Figure 2B, H**), suggesting that the K_V_2.1 expression is similar in these cells.

Further analysis of K_V_2.1 clusters was restricted to those that overlapped with the WGA-mapped plasma membrane. In both, the K_V_2.1_WT_ and K_V_2.1_S590A_ males, the frequency distribution of K_V_2.1 cluster sizes were similar in terms of relative values in all volume bins and could be fit with an exponential decay function (**Figure 2C**). The mean cluster volume in K_V_2.1_WT_ males was 0.07 ± 0.01 μm^3^ (median = 0.07 μm^3^) compared to a mean of 0.07 ± 0.01 μm^3^ (median = 0.06 μm^3^) in K_V_2.1_S590A_ male myocytes. (P = 0.338) (**Figure 2D**). Additionally, total clusters per cell of 551.6 ± 71.1 clusters, (median = 505 clusters) in K_V_2.1_WT_ male myocytes were not significantly different from total clusters per cell of 444.2 ± 33.4 clusters (median = 419 clusters) in K_V_2.1_S590A_ males (P = 0.10) (**Figure 2E)**. The percentage of the membrane occupied by clusters in K_V_2.1_WT_ male myocytes was on average 5.5 ± 0.9% (median = 5.1%), similar to the average in K_V_2.1_S590A_ males of 5.0 ± 1.0% (median = 4.6%) (P = 0.34) (**Figure 2F**).

In sharp contrast to male myocytes, K_V_2.1_S590A_ female myocytes exhibited an increased proportion of smaller clusters as compared to those from K_V_2.1_WT_ females (**Figure 2I**). Accordingly, mean cluster size of K_V_2.1_WT_ in female myocytes was 0.14 ± 0.10 μm^3^ (median = 0.14 μm^3^), significantly larger than mean cluster size of 0.10 ± 0.01 μm^3^ (median = 0.09 μm^3^) (P = 0.007) in K_V_2.1_S590A_ females (**Figure 2J)**. K_V_2.1_WT_ female myocytes had 796 ± 67 clusters per cell, (median = 815 clusters) compared to only 422 ± 34 clusters per cell (median = 415 clusters) in K_V_2.1_S590A_ female myocytes (P < 0.0001) **(Figure 2K)**. Similarly, the percentage of the plasma membrane occupied by K_V_2.1 was higher in K_V_2.1_WT_ female myocytes at 16.7 ± 1.8% (median = 17.7%) in contrast to K_V_2.1_S590A_ female myocytes in which K_V_2.1 clusters occupied on average 7.2 ± 1.0% of the plasma membrane (median = 6.0) (P < 0.0001) (**Figure 2L)**. The significantly lower K_V_2.1 clustering profile in all metrics measured indicates that unlike in males, the S590A mutation decreases channel clustering in female myocytes.

Using the threshold set from our confocal imaging (i.e., macro-clusters are >0.025μm^3^), we quantified the number of macro-clusters expressed in myocytes from K_V_2.1_WT_ and K_V_2.1_S590A_ mice. Around 62% of clusters in K_V_2.1_WT_ male myocytes were identified as macro-clusters, with a similar percentage of 58% observed in samples from males with the K_V_2.1_S590A_ mutation. Approximately 70% of K_V_2.1 clusters in K_V_2.1_WT_ female myocytes were classified as macro-clusters. Remarkably, in K_V_2.1_S590A_ female myocytes, macro-clusters accounted for approximately 49% of the total K_V_2.1 clusters.

We also quantified K_V_2.1 micro-clusters. Although the proportion of micro-clusters was similar in male K_V_2.1_WT_ and K_V_2.1_S590A_ myocytes (38% and 42%, respectively), female K_V_2.1_S590A_ myocytes exhibited a much larger proportion of micro-clusters (51%) compared to myocytes from K_V_2.1_WT_ females (30%). Hence, it can be reasoned that the S590A mutation has a sex-specific effect of reducing the extent of K_V_2.1 macro-clustering in female but not male arterial myocytes without impacting channel expression.

### K_V_2.1 S590 phosphorylation is higher in myocytes from female versus male K_V_2.1_WT_ mice

To investigate the potential role of the S590 phosphorylation site in the sex-specific differences in K_V_2.1 clustering, we conducted immunocytochemistry analyses on arterial myocytes (**Figure 3**). We tested the hypothesis that K_V_2.1_WT_ female myocytes exhibit a higher degree of K_V_2.1 S590 phosphorylation compared to males, which could contribute to the observed sex-specific variations in K_V_2.1 clustering. Accordingly, we utilized a monoclonal antibody (mAb L100/1; (*30*)) specific for K_V_2.1 that is phosphorylated at serine 590 (pS590). **Figure 3A** shows cells double immunolabeled for pS590 (left) and total K_V_2.1 (right) using confocal microscopy.

**Figure 3.**
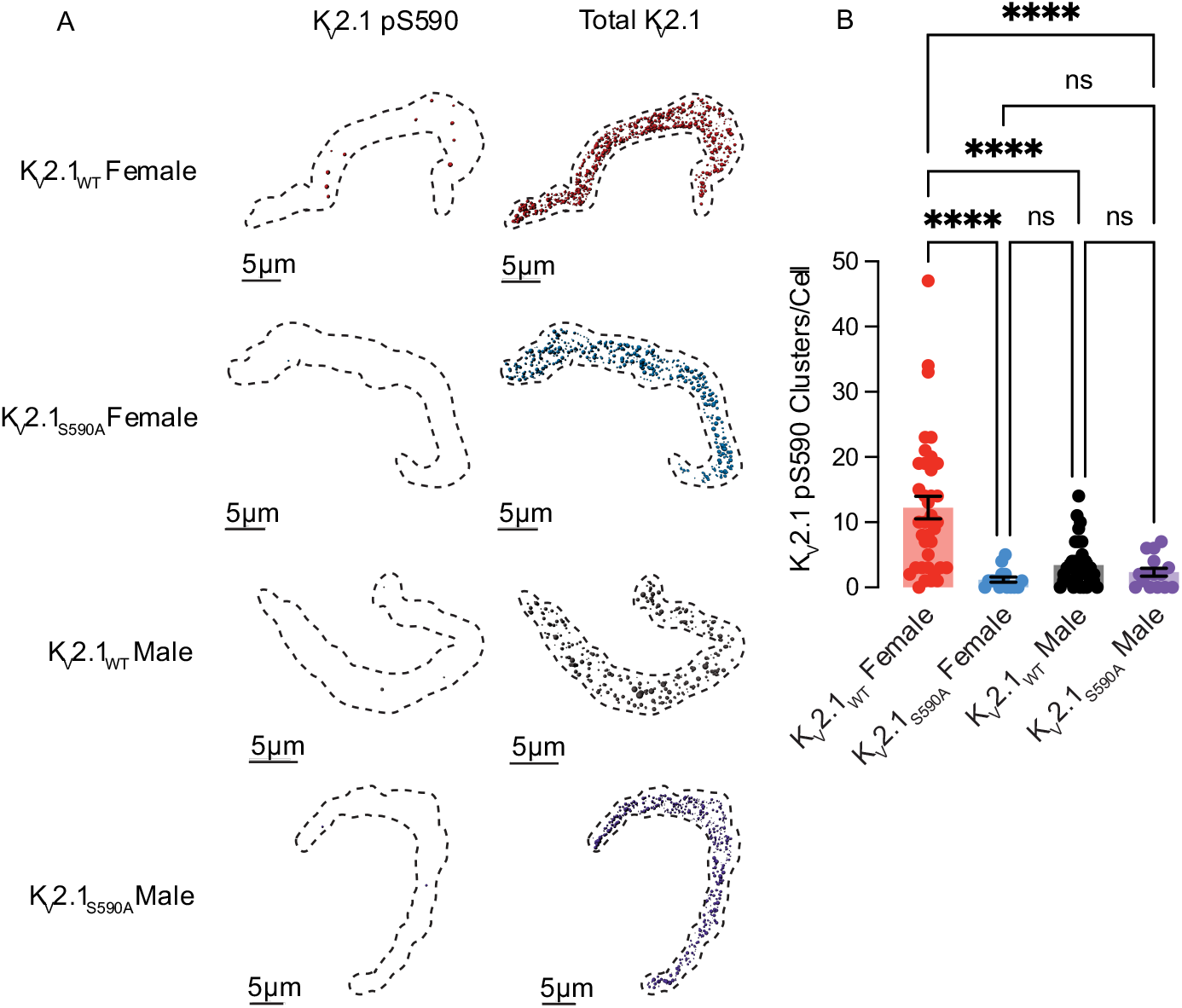
K_V_2.1_WT_ female myocytes exhibit more extensive K_V_2.1 pS590 phosphorylation than those from K_V_2.1_WT_ males. **(A)** Representative maximum projection images of immunolabeling for pS590 K_V_2.1 (left) and total K_V_2.1channel clusters (right) at the surface membrane of K_V_2.1_WT_ female (red), K_V_2.1_S590A_ female (blue), K_V_2.1_WT_ male (black), and K_V_2.1_S590A_ male (purple) cells. (**B**) Summary data of pS590 K_V_2.1 clusters per cell. *P<0.05, **P<0.01, ***P<0.001, ****P<0.0001. Error bars indicate mean ± SEM.

One advantage of our study is that our K_V_2.1_S590A_ mice serve as an ideal negative control. As expected, K_V_2.1_S590A_ males exhibited 2.3 ± 0.6 clusters (median = 2.0 clusters) per cell and K_V_2.1_S590A_ females exhibited 1.2 ± 0.4 clusters (median = 1.0 clusters) per cell confirming the specificity of this mAb for K_V_2.1 phosphorylated at S590. Remarkably, phosphorylated K_V_2.1 clusters were observed in K_V_2.1_WT_ females (**Figure 3B**) exhibiting on average 12.2 ± 1.7 of phosphorylated clusters (median = 10.0 clusters) per cell, whereas K_V_2.1_WT_ males exhibited 3.4 ± 0.6 clusters (median = 2.5 clusters) per cell. Collectively, these findings suggest that basal levels of K_V_2.1 S590 phosphorylation are higher in K_V_2.1_WT_ females, which could account for their increased clustering of K_V_2.1. Furthermore, these data support the notion that K_V_2.1 phosphorylation in K_V_2.1_WT_ male myocytes is constitutively low, making the S590A mutation functionally indistinguishable from non-phosphorylated Kv2.1_WT_ and thus ineffective in altering K_V_2.1 clustering in male myocytes.

### Expression of clustering impaired K_V_2.1_S590A_ does not affect channel activity in arterial myocytes

Three studies, one using Xenopus oocytes(*28*), one using HEK293T cells (*19*) and another from our group using arterial myocytes (*15*) have suggested that the vast majority of K_V_2.1 channels (i.e., 98-99%) expressed in the plasma membrane of these cells are non-conductive. O’Connell *et al(19*) suggested that, at least in HEK293T cells, K_V_2.1 channel activity depends on their density and that channels within large, dense macro-clusters are non-conductive. A testable hypothesis raised by these data is that a larger fraction of K_V_2.1_S590A_ should be conductive and hence the amplitude of K_V_2.1 currents in native arterial myocytes should differ between cells from K_V_2.1_WT_ and K_V_2.1_S590A_ mice (**Figure 4**).

**Figure 4.**
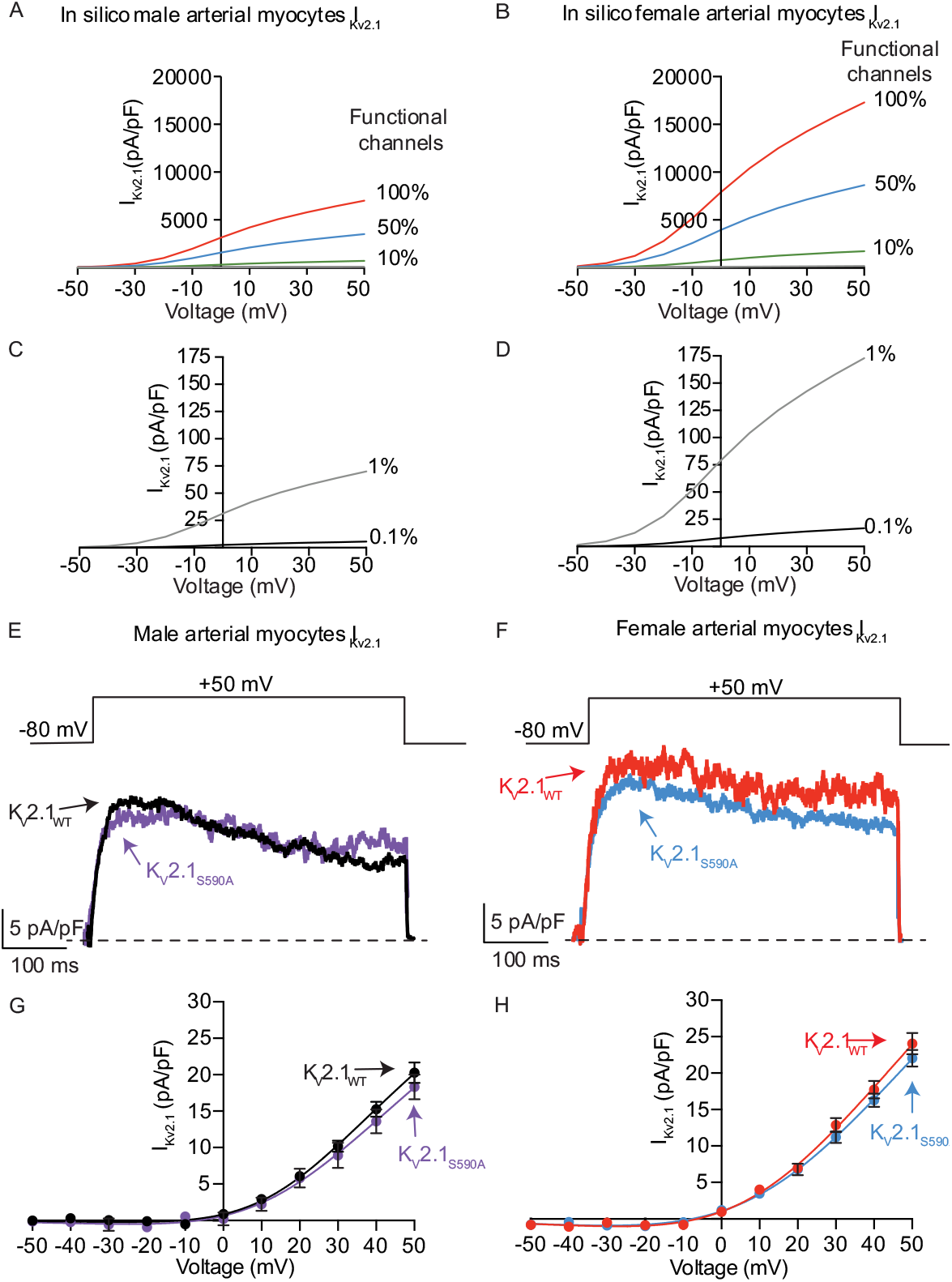
Expression of clustering impaired K_V_2.1_S590A_ does not affect K_V_2.1 channel activity in arterial myocytes. Computationally modeled I_Kv2.1_ in male (**A**) and female (**B**) myocytes assuming 100% (red), 50% (blue) or 10% (green) of K_V_2.1 channels present in the plasma membrane are functional. Computationally modeled I_Kv2.1_ in male (**C**) and female (**D**) myocytes assuming 1% (gray) and 0.1% (black) K_V_2.1 channels are functional. (**E**) Representative I_Kv2.1_ traces at +50 mV from K_V_2.1_WT_ male (black) and K_V_2.1_S590A_ male (purple) arterial myocytes. (**F**) Representative I_Kv2.1_ traces at +50 mV from K_V_2.1_WT_ female (red) and K_V_2.1_S590A_ (blue) arterial myocytes. I_KV2.1_ traces were obtained by subtracting currents after the application of RY785 from control I_K_ traces. Voltage dependence of I_Kv2.1_ in K_V_2.1_WT_ and K_V_2.1_S590A_ male (**G**, black and purple) and K_V_2.1_WT_ and K_V_2.1_S590A_ female (**H**, red and blue) myocytes. Error bars indicate mean ± SEM.

We tested this hypothesis using a multipronged approach. First, we used a mathematical modeling approach(*31*) to determine the predicted changes in macroscopic K_V_2.1 currents with varied levels of functional channels (i.e., 0.1, 1, 10, 50, or 100%) in male and female arterial myocytes (**Figure 4A-D**). The rationale for this analysis is that it provides a set of potential outcomes that can provide insights into the degree of K_V_2.1 declustering in K_V_2.1_S590A_ myocytes. This model incorporated data (e.g., voltage-dependencies and number of channels in the sarcolemma) from O’Dwyer et al(*15*).

As shown in **Figure 4A**, the model predicts that with 100%, 50%, or 10% functional K_V_2.1 channels in male myocytes would produce current densities at +50 mV of 7,006, 3,503, and 701 pA/pF, respectively. By contrast, at the same voltage, the *in silico* female arterial myocytes produce current densities of 17,293, 8,646, and 1,729 pA/pF with 100%, 50%, or 10% functional K_V_2.1 channels (**Figure 4B**). We also simulated the current-voltage relationships in male (**Figure 4C**) and female (**Figure 4D**) myocytes assuming 1% and 0.1% of K_V_2.1 channels are conductive, which are more within the range with previous experimental results in heterologous systems (*19, 28*) and native cells (*15*). The magnitude of *in silico* K_V_2.1 current densities with 1% or 0.1% functional channels was 70.1 and 5.57 pA/pF in male myocytes and 173 and 16.7 pA/pF in female myocytes.

Next, we recorded voltage-gated K^+^ (K_V_) currents in male and female K_V_2.1_WT_ and K_V_2.1_S590A_ arterial myocytes in response to 500 ms depolarizations to voltages between -50 and +50 mV before and after applying the K_V_2.1 blocker RY785 (1 μM) (*32, 33*). This compound decreases K_V_2.1 currents by blocking the pore of these channels (*32*) rather than by immobilizing their voltage sensor, as stromatoxin does (*34*). As a first step in these experiments, we tested the specificity of the RY785 by recording Kv currents before and after the application of this molecule in male and female K_V_2.1_WT_ and K_V_2.1 null (K_V_2.1^-/-^) myocytes (**Supplemental Figure 2A-D)**. Notably, application of 1 μM RY785 decreased the amplitude of K^+^ currents in K_V_2.1_WT_ but not in K_V_2.1^-/-^ myocytes of either sex. This indicates that RY785 is a specific blocker of K_V_2.1 channels in arterial myocytes.

Having completed these critical control experiments, we recorded K_V_ currents from K_V_2.1_WT_ and K_V_2.1_S590A_ myocytes. We noted that the amplitude of the composite K currents were similar in myocytes from K_V_2.1_S590A_ mice compared to myocytes from sex-matched K_V_2.1_WT_ littermates (**Supplemental Figure 2E**. Importantly, for both sexes, RY785-sensitive K_V_2.1 currents were also similar in male (**Figure 4E, G**) and female (**Figure 4F, H**) K_V_2.1_WT_ and K_V_2.1_S590A_ myocytes. Indeed, a comparison of the experimental and in silico amplitudes of the macroscopic K_V_2.1 currents suggests that less than 1% of the channels are functional in myocytes from both male and female K_V_2.1_WT_ and K_V_2.1_S590A_ mice. When taken together with our analyses of Kv2.1 clustering detailed above, these findings suggest that in arterial myocytes K_V_2.1 channel activity is not determined by the extent and nature of its clustering.

### The K_V_2.1_S590A_ mutation diminishes K_V_2.1 and Ca_V_1.2 interactions in female myocytes

We used the proximity ligation assay (PLA) to interrogate the impact of the K_V_2.1_S590A_ mutation on protein-protein interactions, at a resolution of approximately 40 nm (*35, 36*) (**Supplemental Figure 3**). We first evaluated K_V_2.1-K_V_2.1 interactions within isolated mesenteric smooth muscle cells by using two different antibodies directed against different epitopes in the K_V_2.1 cytoplasmic C-terminus. In this case both intra- and inter-molecular proximity of the two epitopes would yield a PLA signal. Inter-molecular interactions could be visualized as “puncta”, and we hypothesized that more puncta would be exhibited in cells where K_V_2.1 was more clustered since there would be increased proximity of epitopes due to intermolecular interactions, allowing for more PLA reactions to occur. Confocal images of K_V_2.1_WT_ and K_V_2.1_S590A_ myocytes subjected to PLA show that puncta of K_V_2.1-K_V_2.1 PLA signals were randomly distributed throughout the cell, and that PLA signal could be detected in all cells consistent with our confocal data above showing that K_V_2.1 micro-clustering still occurs in mutant myocytes of both sexes (**Supplemental Figure 3A**). The density of PLA puncta of 0.035 ± 0.003 puncta/μm^2^ (median = 0.035 μm^2^) in K_V_2.1_WT_ males was similar to 0.030 ± 0.003 puncta/μm^2^ (median = 0.028 μm^2^) in K_V_2.1_S590A_ male mice (P = 0.148) (**Supplemental Figure 3B**). Consistent with the confocal imaging analysis, the density of K_V_2.1-K_V_2.1 PLA puncta was greater in K_V_2.1_WT_ females, with an average of 0.153 ± 0.011 puncta/μm^2^ (median = 0.162 μm^2^), compared to 0.046 ± 0.003 puncta/μm^2^ (median = 0.037 μm^2^) in K_V_2.1_S590A_ females (P < 0.0001) (**Supplemental Figure 3C**), suggesting that the K_V_2.1_S590A_ mutation reduces the level of K_V_2.1 clustering in female myocytes.

Previous work from our group has shown that K_V_2.1 expression promotes Ca_V_1.2 clustering and activity in neurons (*16*) and arterial myocytes (*15*). Following from this and the data above, we hypothesize that in arterial myocytes K_V_2.1 plays a sex-specific structural role as an organizer to bring Ca_V_1.2 channels together in female but not male myocytes. We again used PLA to test the hypothesis that the declustering of K_V_2.1 channels in female but not male myocytes from K_V_2.1_S590A_ mice would decrease K_V_2.1-Ca_V_1.2 channel interactions in a sex-specific manner. Representative images of K_V_2.1-Ca_V_1.2 PLA puncta show randomly distributed interactions across the cell (**Supplemental Figure 3D**).

Quantification showed that K_V_2.1-Ca_V_1.2 puncta density was unchanged in males between K_V_2.1_WT_ male myocytes with a mean of 0.022 ± 0.002 puncta/μm^2^ (median = 0.016 puncta/μm^2^) and K_V_2.1_S590A_ male myocytes with a mean of 0.018 ± 0.002 puncta/μm^2^ (median = 0.016 puncta/μm^2^; P = 0.217) (**Supplemental Figure 3E**). However, K_V_2.1-Ca_V_1.2 interactions decreased in female K_V_2.1_S590A_ myocytes with a mean of 0.030 ± 0.004 puncta/μm^2^ (median = 0.026 puncta/μm^2^) compared to K_V_2.1_WT_ female myocytes with a mean of 0.044 ± 0.004 puncta/μm^2^ (median = 0.026 puncta/μm^2^; P = 0.013) (**Supplemental Figure 3F**). These data further support a sex-specific structural role for K_V_2.1 channels, facilitating Ca_V_1.2-Ca_V_1.2 clustering.

### Female myocytes expressing K_V_2.1_S590A_ have reduced macroscopic Ca_V_1.2 currents

We recorded macroscopic Ca_V_1.2 currents (I_Ca_) from male and female K_V_2.1_WT_ and K_V_2.1_S590A_ arterial myocytes (**Figure 5A-D**). I_Ca_ was activated by applying 300 ms voltage step depolarizations from a holding potential of -80 to +60 mV. We show I_Ca_ traces recorded during a depolarization to 0 mV from representative male (**Figure 5A**) and female (**Figure 5B**) K_V_2.1_WT_ and K_V_2.1_S590A_ arterial myocytes. Note that the amplitude and kinetics of I_Ca_ in these male K_V_2.1_WT_ and K_V_2.1_S590A_ arterial myocytes were similar. By contrast, we found that peak I_Ca_ was smaller in female Kv2.1_S590A_ myocytes compared to those in K_V_2.1_WT_ cells. In **Figure 5C** and **D**, we show the voltage dependence of the amplitude of I_Ca_ from all the cells examined over a wider range of membrane potentials. This analysis shows that the amplitude of I_Ca_ is similar in K_V_2.1_WT_ and K_V_2.1_S590A_ male myocytes at all voltages examined. However, in female myocytes, I_Ca_ was smaller in K_V_2.1_S590A_ than in K_V_2.1_WT_ at all voltages examined. Indeed, at 0 mV, I_Ca_ amplitude in K_V_2.1_S590A_ cells was approximately 50% of that of WT females.

**Figure 5.**
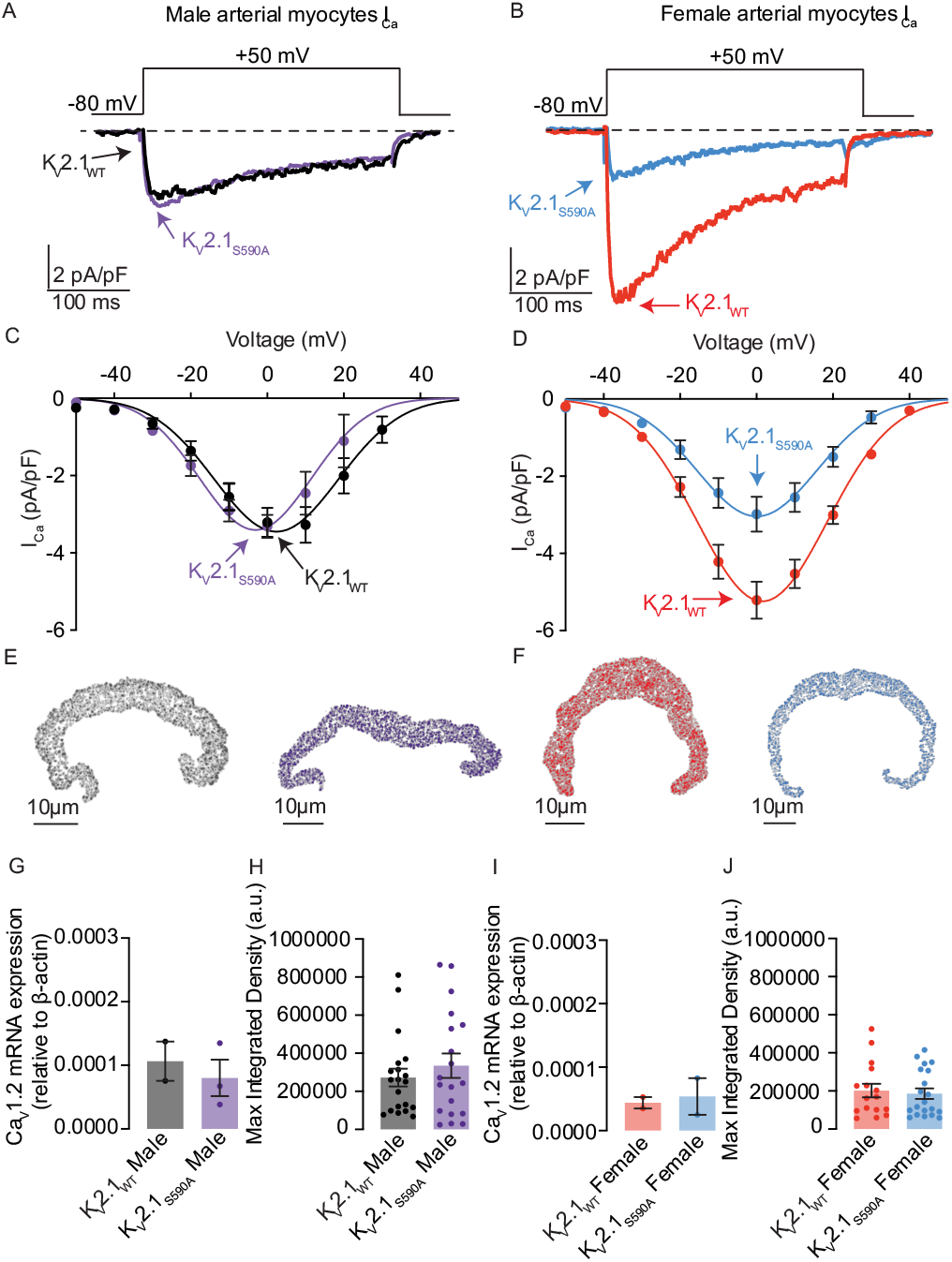
I_Ca_ is reduced in K_V_2.1_S590A_ female myocytes but unaffected in male arterial myocytes. I_Ca_ records (0 mV) from representative K_V_2.1_WT_ and K_V_2.1_S590A_ male (**A**) and K_V_2.1_WT_ and K_V_2.1_S590A_ female (**B**) myocytes. Voltage dependence of I_Ca_ from male (**C**) and female (**D**) myocytes at membrane potentials ranging from -50 to +50 mV. (**E**,**F**) Representative images of immunolabeled Ca_V_1.2 in myocytes from K_V_2.1_WT_ male (**5E**, black), K_V_2.1_S590A_ male (**5E**, purple), K_V_2.1_WT_ female (**5F**, red), and K_V_2.1_S590A_ female (**5F**, blue) mice. Summary data from real-time quantitative PCR experiments of Ca_V_1.2 mRNA expression relative to β-actin in male (**G**) and female (**I**) myocytes. Quantification of immunofluorescence of labeled Ca_V_1.2ɑ subunit in male (**H**) and female (**J**) myocytes. Error bars indicate mean ± SEM.

Next, we determined the level of expression of Ca_V_1.2 protein in male and female K_V_2.1_S590A_ and K_V_2.1_WT_ vessels using immunocytochemistry (**Figure 5E, F, H, J**) and RT-PCR approaches (**Figure 5G and I**). Our analysis suggests that total Ca_V_1.2 protein and mRNA expression is similar in male and female K_V_2.1_S590A_ and K_V_2.1_WT_ vessels. This suggests that the smaller I_Ca_ in female K_V_2.1_S590A_ than K_V_2.1_WT_ myocytes is not likely due to lower Ca_V_1.2 expression in these cells.

### Declustering K_V_2.1 in myocytes with the Kv2.1_S590A_ mutation decreases Ca_V_1.2 cluster sizes in female but not male arterial myocytes

In a previous study (*15*), we suggested a model that differences in I_Ca_ amplitude between female and male arterial myocytes were due to sex-specific differences in K_V_2.1-mediated Ca_V_1.2 clustering that impacted the probability of cooperative gating of these channels. Our data above show differences in I_Ca_ amplitude between female K_V_2.1_WT_ and K_V_2.1_S590A_ myocytes in the absence of differences in Ca_V_1.2 expression levels. Thus, we investigated whether K_V_2.1_S590A_ expression altered Ca_V_1.2 channel clustering in a sex-specific manner using ground state depletion (GSD) super-resolution microscopy (**Figure 6**).

**Figure 6.**
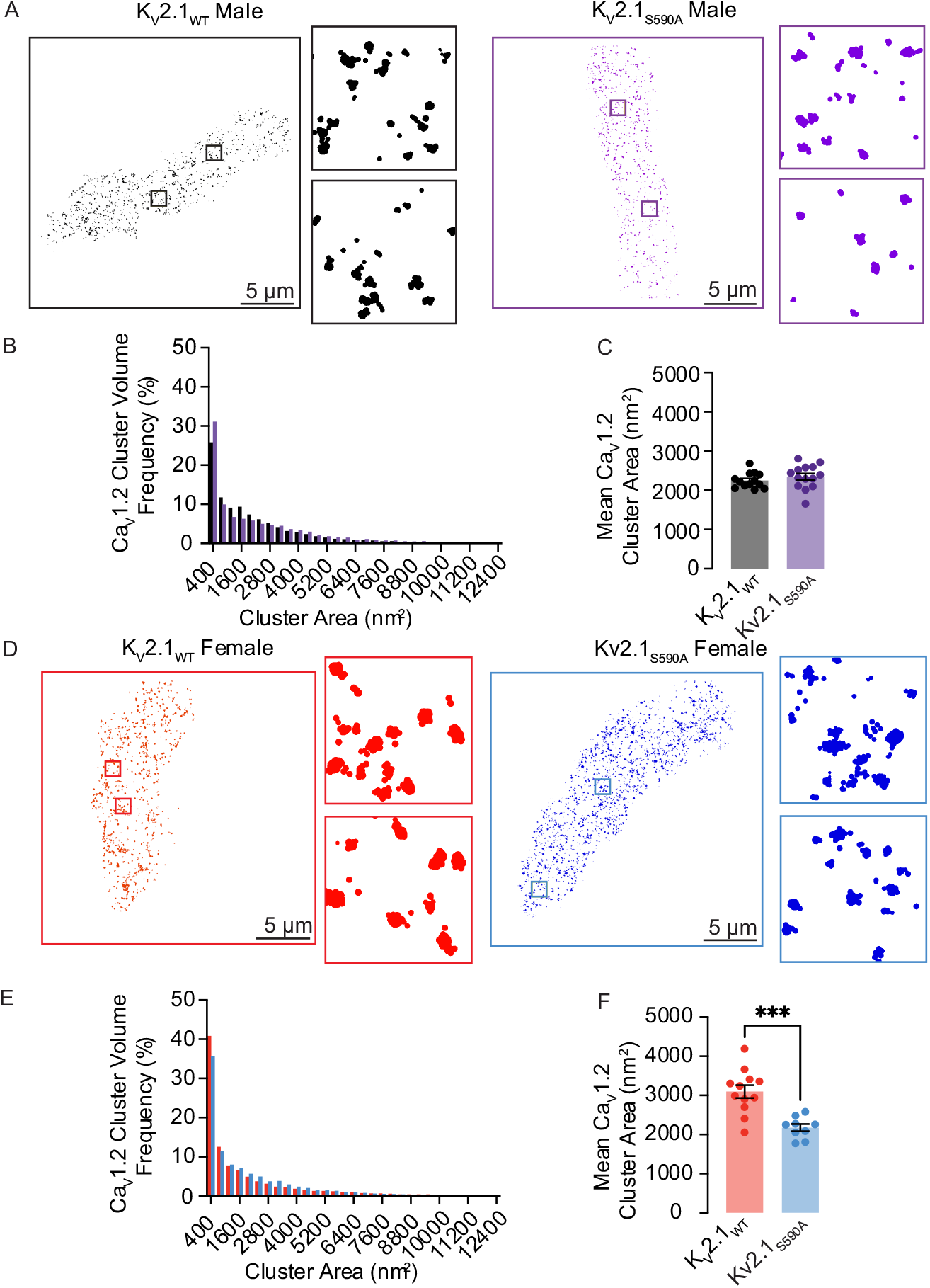
Ca_V_1.2 cluster sizes are decreased in myocytes from female but not male K_V_2.1_S590A_ mice. (**A**) Representative super-resolution GSD images of immunolabeled Ca_V_1.2 labeled channels in K_V_2.1_WT_ (left) and K_V_2.1_S590A_ (right) male myocytes. Insets show 4 μm^2^ regions of interest. (**B**) Relative frequency as a percentage of Ca_V_1.2 cluster areas of K_V_2.1_WT_ (black) and K_V_2.1_S590A_ (purple) in male myocytes. (**C**) Summary plot of mean Ca_V_1.2 cluster areas in male myocytes (**D**) Representative super-resolution GSD microscopy images of immunolabeled Ca_V_1.2 labeled channels in K_V_2.1_WT_ (left) and K_V_2.1_S590A_ (right) female myocytes. (**E**) Relative frequency as a percentage of Ca_V_1.2 cluster areas of K_V_2.1_WT_ (red) and K_V_2.1_S590A_ (blue) female myocytes. (**F**) Summary plot of mean Ca_V_1.2 cluster areas in female myocytes. *P < 0.05, **P < 0.01, ***P < 0.001. Error bars indicate mean ± SEM.

We show ground-state depletion run in TIRF mode super-resolution images from representative male (**Figure 6A**) and female (**Figure 6D**) myocytes from K_V_2.1_WT_ and K_V_2.1_S590A_ mice. The insets show expanded views of two regions of interest (1 μm^2^) within each cell image. Our TIRF images show that Ca_V_1.2 clusters of various sizes are expressed throughout these cells. The frequency distribution of Ca_V_1.2 cluster areas in K_V_2.1_WT_ and K_V_2.1_S586A_ male and females could both be fit with an exponential function (**Figure 6B, E)**.

The mean area of Ca_V_1.2 clusters in male K_V_2.1_WT_ of 2259 ± 55 nm^2^ (median = 2219 nm^2^) was similar to the K_V_2.1_S590A_ male mean of 2345 ± 82 nm^2^ (median = 2354 nm^2^) (P = 0.173) (**Figure 6C**), suggesting that declustering K_V_2.1 in male myocytes does not affect Ca_V_1.2 channel clustering. However, Ca_V_1.2 cluster sizes were significantly smaller in K_V_2.1_S590A_ female myocytes with a mean area of 2381 ± 91 nm^2^ (median = 2251 nm^2^) compared to K_V_2.1_WT_ female myocytes whose mean area was 3098 ± 164 nm^2^ (median = 3117 nm^2^) (P = 0.0001). Taken together with our electrophysiological data, our findings suggest that the clustering and activity of Ca_V_1.2 channels is modulated by the degree of K_V_2.1 clustering.

As shown in **Supplemental Figure 4A and B**, our stochastic self-assembly model effectively reproduced the steady-state size distributions that we measured for Ca_V_1.2 clustering in K_V_2.1_WT_ and K_V_2.1_S586A_ arterial myocytes. The parameters used in the model are summarized in **Supplemental Figure 4C**. These *in silico* data suggest that Ca_V_1.2 clusters in K_V_2.1_S590A_ female myocytes have a higher probability of growth (i.e., *P*_*g*_) than those in female K_V_2.1_WT_ arterial myocytes.

### K_V_2.1_S586A_ reduces Ca_V_1.2-Ca_V_1.2 channel interactions

Having previously shown that K_V_2.1 enhances Ca_V_1.2-Ca_V_1.2 channel interactions in arterial myocytes (*15*), we set out to determine if a decrease in K_V_2.1 clustering would lead to a reduction in Ca_V_1.2-Ca_V_1.2 interactions. We utilized a split-Venus fluorescent protein system to visualize Ca_V_1.2 channels. This system involves fusing Ca_V_1.2 channels with either the N-terminal fragment (Ca_V_1.2-VN) or the C-terminal fragment (Ca_V_1.2-VC) of Venus protein. Individually, neither Ca_V_1.2-VN nor Ca_V_1.2-VC emits fluorescence. However, when brought into close enough proximity for interaction to occur, they can reconstitute a full fluorescence emitting Venus protein. Thus, the split-Venus fluorescence can be utilized to indicate spontaneous interactions between neighboring Ca_V_1.2 channels. Accordingly, we compared the split-Venus fluorescence in HEK293T cells expressing Ca_V_1.2-VN and Ca_V_1.2-VC and co-expressing either K_V_2.1_WT_ or K_V_2.1_S586A_ (**Figure 7**). The voltage protocols used for these experiments are similar to those used in two recent studies (*10, 15*) and are described in detail in the Methods section of this paper.

**Figure 7.**
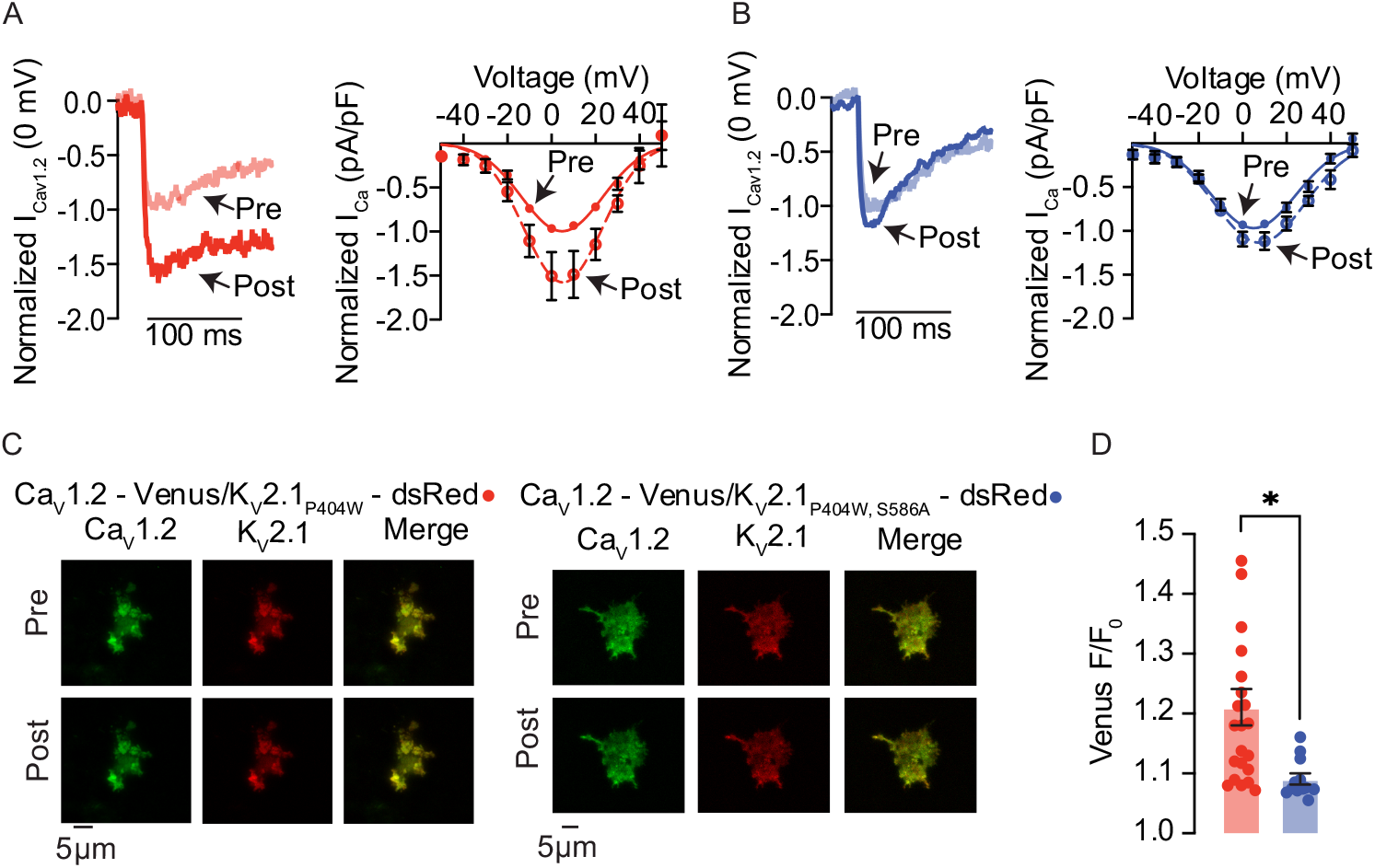
Ca_V_1.2-Ca_V_1.2 interactions are decreased in cells expressing K_V_2.1_S586A_. (**A**, left) Representative currents measured at 0 mV from pre-(opaque red) and post-(red) conditioning protocol from HEK293T cells expressing Ca_V_1.2-VN, Ca_V_1.2-VC, and DsRed-K_V_2.1_P404W_. (**A**, right) Normalized (to peak current in pre-conditioning protocol) pre- and post-IV relationships from HEK293T cells expressing Ca_V_1.2-VN, Ca_V_1.2-VC, and DsRed-K_V_2.1_P404W_. (**B**, left) Representative currents measured at 0 mV from pre-(opaque blue) and post-(blue) conditioning protocol from HEK293T cells expressing Ca_V_1.2-VN, Ca_V_1.2-VC, and DsRed-K_V_2.1_P404W,S586A_. (**B**, right) Normalized pre- and post-IV relationships from HEK293T cells expressing Ca_V_1.2-VN, Ca_V_1.2-VC, and DsRed-K_V_2.1_P404W,S586A_. (**C**) Representative TIRF images of Venus fluorescence reconstitution in HEK293T cells from cells transfected with Ca_V_1.2-VC, Ca_V_1.2-VC and DsRed-K_V_2.1_P404W_ (left) or Ca_V_1.2-VC, Ca_V_1.2-VC and DsRed-K_V_2.1_P404W,S586A_ (right). Pre- and post-conditioning Ca_V_1.2-Venus (green), K_V_2.1_P404W_ or K_V_2._1P404W,S856A_ (red), and the merge of the two channels are presented. (**D**) Summary of Ca_V_1.2-Venus fluorescence (F/F_0_). *P < 0.05. Error bars indicate mean ± SEM.

We first transfected HEK293T cells with Ca_V_1.2-VC, Ca_V_1.2-VN, and the non-conducting but clustering competent rat K_V_2.1_P404W_ channel(*37*) tagged with red-shifted fluorescent protein dsRed. The P404W mutation confers a non-conductive K_V_2.1 phenotype, allowing us to study the structural clustering role of K_V_2.1 without masking of the Ca^2+^ currents by K^+^. We found that I_Ca_ was larger at most membrane potentials in our post-conditioning IV protocol with a peak I_Ca_ at 0 mV showing an increase by about 51% (**Figure 7A**). Representative TIRF images are provided before and after conditioning protocol (**Figure 7C**). Note the appearance of co-clusters of Ca_V_1.2 (green) and K_V_2.1 (red). Our data show that Venus fluorescence increased by approximately 21% with stimulation from basal to post-conditioning steps suggesting an increase in K_V_2.1 dependent Ca_V_1.2-Ca_V_1.2 interactions (**Figure 7D**). This supports previously published results(*15, 17*). To test the role of declustering K_V_2.1 with the S586A point mutation on Ca_V_1.2 channel interactions, we co-expressed Ca_V_1.2-VN, Ca_V_1.2-VC and DsRed-K_V_2.1_P404W,S586A_ in HEK293T cells and repeated the above protocol. We found that I_Ca_ did exhibit a small increase of about 9.4% between pre- and post-conditional protocols (**Figure 7B**). In representative TIRF footprints, we show that Ca_V_1.2-VN, Ca_V_1.2-VC and K_V_2.1_P404W_,_S586A_ transfected cells expressed more diffusely clustered Ca_V_1.2 and K_V_2.1 channels, visually confirming that K_V_2.1 is declustered (**Figure 7C**). Furthermore, Venus fluorescence with K_V_2.1_P404W,S586A_ expression increased by about 9%, a level similar to what was previously published(*15, 17*) with Ca_V_1.2-VN and Ca_V_1.2 VC alone (**Figure 7D)**. We propose this small increase is due to the intrinsic ability of Ca_V_1.2 channels to interact with one another. However, this increase in Venus fluorescence was significantly lower than that seen in K_V_2.1_P404W_ transfected cells (**Figure 7D)**. Together these data further support the structural role K_V_2.1 channels play in modulating Ca_V_1.2-Ca_V_1.2 interactions and activity.

### The activity of Ca_V_1.2 channels is reduced in K_V_2.1_S590A_ female but not male arterial myocytes

We next examined whether variations in the activity of Ca_V_1.2 channels could explain the differences in I_Ca_ observed in myocytes from K_V_2.1_WT_ and K_V_2.1_S590A_ male and female mice. Ca_V_1.2 channel activity was determined by recording Ca_V_1.2 sparklets using TIRF microscopy as previously described (*6, 7, 12, 17, 38-40*) (**Figure 8**).TIRF microscopy of near-plasma membrane intracellular Ca^2+^ levels provides a powerful tool for recording Ca^2+^ entry via individual or small clusters of Ca_V_1.2 channels, as it enables the activity of individual channels to be recorded from a relatively large membrane area allowing for the identification of discrete sarcolemma signaling domains. In this analysis, Ca_V_1.2 sparklet activity is expressed as nP_s_, where n is the number of quantal levels reached by the sparklet site and P_s_ is the probability of sparklet occurrence.

**Figure 8.**
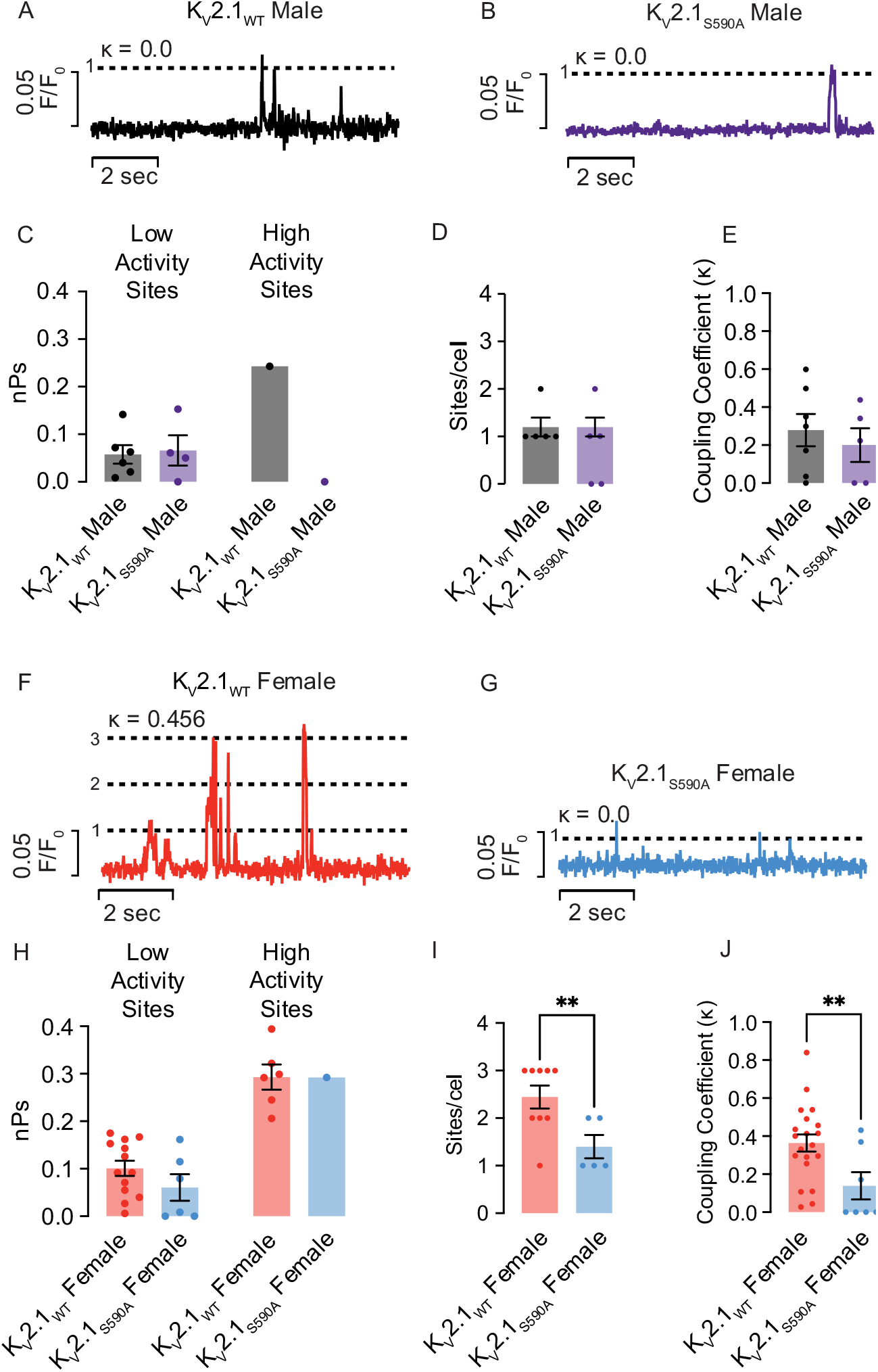
Activity of Ca_V_1.2 channels is reduced in K_V_2.1_S586A_ female but not male arterial myocytes. Representative sparklet traces from K_V_2.1_WT_ male (**A**), K_V_2.1_S590A_ male (**B**), K_V_2.1_WT_ female (**F**), and K_V_2.1_S590A_ female (**G**) myocytes. κ values are shown above each trace. nPs values from low activity sites (left) and high activity sites (right) in male (**C**) and female (**H**) myocytes. Coupling coefficient values (κ) from male (**D**) and female (**I**) myocytes. Sparklet sites per cell from male myocytes (**E**) and female (**J**) myocytes. *P < 0.05, **P < 0.01. Error bars indicate mean ± SEM.

As previously reported (*38*), detailed analysis of Ca_V_1.2 sparklets sites revealed heterogeneity in activity at different sites. Therefore, Ca_V_1.2 sparklets sites were separated into low and high activity sites, using an nP_s_ cutoff of 0.2.

Representative Ca_V_1.2 sparklet traces are provided (**Figure 8 A, B, F, G**) from low activity sparklet sites. Of note, the majority of the sparklet activity that occurs in male myocytes is produced by a signal that corresponds to a single channel opening (one quantal unit) (**Figure 8A, B**). The strength of the coupled gating is denoted by the κ value, and in these traces, the κ values are close to or equal to 0, indicating no or weak coupling between the channels. In contrast, the K_V_2.1_WT_ female trace (**Figure 8F)** from a low activity site exhibited coordinated multi-channel openings, of up to 3 channels with a κ value of 0.466. Interestingly, the activity of sparklet sites from K_V_2.1_S590A_ female myocytes (**Figure 8G)** were similar to those of K_V_2.1_WT_ and K_V_2.1_S590A_ male myocytes (**Figure 7A, B)**, exhibiting mostly single channel openings and few coupled gating events.

We found that in low activity sparklet sites, the average nP_s_ was not significantly different between K_V_2.1_WT_ and K_V_2.1_S590A_ male myocytes (**Figure 8C**). K_V_2.1_WT_ sparklet sites had an average nP_s_ of 0.06 ± 0.02 (median = 0.05) compared to K_V_2.1_S590A_ where the nP_s_ average was 0.07 ± 0.03 (median = 0.06) (P = 0.41). In male myocytes of either genotype, we rarely observed cells exhibiting high activity sparklet sites, except for a single site in a K_V_2.1_WT_ male myocyte (**Figure 8C**). Furthermore, we did not observe a difference in the number of Ca_V_1.2 sparklet sites between K_V_2.1_WT_ and K_V_2.1_S590A_ male myocytes, with most cells exhibiting just one site (**Figure 8D)**.

Similarly, when we compared nP_s_ in low activity sites in K_V_2.1_WT_ and K_V_2.1_S590A_ female myocytes, we could not discern a difference in their average nP_s_ values (**Figure 7H**). K_V_2.1_WT_ female mean nPs was 0.10 ± 0.02 (median = 0.09), which was similar to 0.06 ± 0.03 (median = 0.04) in K_V_2.1_S590A_ female myocytes (P = 0.10). High activity sites averaged an nP_s_ of 0.29 ± 0.03 (median = 0.30) in K_V_2.1_WT_ cells and a mean of 0.29 ± 0.01 (median = 0.29) in K_V_2.1_S590A_ female cells (P = 0.50) (**Figure 7H**). However, K_V_2.1_WT_ female myocytes exhibited 2.44 ± 0.24 (median = 3.0) Ca_V_1.2 sparklet sites per cell, significantly higher than 1.40 ± 0.25 (median = 1.0) Ca_V_1.2 sparklet sites in K_V_2.1_S590A_ female myocytes (P = 0.008) suggesting decreased Ca_V_1.2 channel activity with S590A mutation (**Figure 7I**).

Previous work (*40*) showed that Ca_V_1.2 sparklet sites appeared to arise from the simultaneous opening and/or closing of multiple channels suggesting that small groups of channels may be functioning cooperatively. To examine such coupling, we employed a coupled Markov chain model to determine the coupling coefficient (κ) among Ca_V_1.2 channels at Ca^2+^ sparklet sites. The κ value ranges from 0 for channels that gate independently to 1 for channels that are tightly coupled and open and close simultaneously. A detailed description of this model is provided in the expanded Methods section.

Using this analysis, we found that the average κ value of 0.28 ± 0.09 (median = 0.30) in K_V_2.1_WT_ male myocytes was not significantly different from 0.20 ± 0.09 (median = 0.22) in K_V_2.1_S590A_ male myocytes (P = 0.27) (**Figure 8E**). However, the average κ value of 0.36 ± 0.05 (median = 0.38) in female K_V_2.1_WT_ myocytes, was significantly higher than 0.14 ± 0.07 (median = 0) (P = 0.0082) in female K_V_2.1_S590A_ myocytes, suggesting more coupled events (**Figure 8J**). Taken together, these data indicate increased Ca_V_1.2 channel activity and coupled gating in myocytes from K_V_2.1_WT_ females compared to those with the K_V_2.1_S590A_ mutation, suggesting that clustering of K_V_2.1 modulates Ca_V_1.2 channel activity.

## Discussion

In this study, we show that arterial smooth muscle cells from mice expressing a gene-edited point mutation of the K_V_2.1 channel that selectively eliminates its characteristic macro-clustered localization have properties remarkably like those from K_V_2.1 knock-out mice. This leads us to formulate a new model in which K_V_2.1 expression — *by itself* — is not sufficient for this channel to exert its structural functions on modulating Ca_v_1.2 clustering and activity, but rather depends on K_V_2.1 channel’s capacity to form macro-clusters. Notably, the presence of K_V_2.1 macro-clusters in female, but not male myocytes underlie sex-specific differences in Ca^2+^ influx via Ca_V_1.2 channels in arterial smooth muscle. Our data suggest a new paradigm whereby the clustering of ion channels underlies their physiological functions, independent of their ability to conduct ions.

Analysis of super-resolution images indicates that clustering of K_V_2.1 and Ca_V_1.2 channels is random and hence does not involve an active process. This stochastic self-assembly mechanism leads to micro- and macro-clusters of varying sizes that represent the default organization of K_V_2.1 and Ca_V_1 channels expressed endogenously in neurons and smooth muscle cells or exogenously in heterologous cells (*9*). Furthermore, we found that K_V_2.1 macro-clusters are composed of groups of micro-clusters. This is consistent with a recent study showing that in developing neurons K_V_2.1 macro-clusters formed from the coalescence of numerous micro-clusters (*41*) and suggests that the organization of K_V_2.1 clusters is hierarchical.

An important finding in this study is that K_V_2.1 clustering is more prominent in female than in male arterial myocytes, with female myocytes expressing a larger proportion of macro-clusters. In this context, the development of the K_V_2.1_S590A_ mouse allowed us to investigate the separatable structural clustering and ion conducting roles of this channel. We found that expression K_V_2.1_S590A_ nearly eliminated macro-clustering in female myocytes but had no impact on K_V_2.1 micro-clusters in cells from male or female myocytes. Because the S590A mutation eliminated a phosphorylation site in the PRC domain that causes macro-clustering, these findings suggested that the potential mechanism of these sex-specific differences in K_V_2.1 clustering was differential phosphorylation of this specific serine in male and female myocytes.

Indeed, K_V_2.1 phosphorylation and macro-clustering is regulated by a myriad of protein kinases such as CDK5 and protein phosphatases such as calcineurin (*42*). These kinases and phosphatases work as a rheostatic mechanism to regulate the phosphorylation status of K_V_2.1 based on physiological demands (*26, 27, 42*). Accordingly, we found that that the phosphorylation state of K_V_2.1 in arterial myocytes differs between the two sexes, specifically, that K_V_2.1 in male myocytes is phosphorylated to a much lower degree.

It is intriguing to consider the potential clustering mechanisms that are impacted by inhibiting phosphorylation at the 590/586 specific site of the PRC domain by the serine to alanine point mutation. One hypothesis is that VAP proteins act to modulate the probability of macro-cluster formation. Studies show that K_V_2.1 clusters are expressed at sites where the endo/sarcoplasmic reticulum is brought into close juxtaposition to the plasma membrane (ER/SR-PM junctions) (*41*) and this interaction and accumulation of channels relies on the tethering of K_V_2.1 to VAP proteins (*21, 43*). The transmembrane ER/SR VAP proteins (VAPA and VAPB) interact with the phosphorylated K_v_2.1 PRC domain and have been proposed to function to increase the local concentration of K_V_2.1 channels at ER/SR-PM junctions resulting in K_V_2.1 macro-clustering.

Consistent with this, Kirmiz et al., (*21*) found that knock-out of VAPA in RAW664.7 macrophage cells resulted in a decrease in K_V_2.1 channel clustering. Knockdown of endogenous VAP proteins similarly impaired clustering of K_V_2.1 heterologously expressed in HEK293T cells (*43*). Interestingly, the model proposed in these prior papers (*21, 43*) suggested that the phosphorylated PRC domain is necessary and sufficient for macro-clustering of K_V_2 channels. This is consistent with prior studies showing that mutations disrupting or eliminating the PRC domain (*22, 37, 43*) or treatments that impact K_V_2.1 phosphorylation (*26, 27, 42*) impact K_V_2.1 clustering. It is presumed that the phosphorylation of multiple serine residues, including S590, within the PRC domain provide the negative charges needed to generate a functional VAP-binding FFAT — two phenylalanines in an acidic tract — motif, as has been shown for numerous other proteins that exhibit phosphorylation-dependent binding to VAPs (*44*).Therefore, one possible mechanism for the decrease in macro-clustering in the S590A mutant is the inability of VAP proteins to recognize the PRC domain of mutated channels preventing cluster growth. The similarity in the patterns of cluster sizes and densities between HEK293T cells and arterial myocytes of both WT and S590A channels is noteworthy, indicating the possibility of a shared set of mechanisms. Further research will be necessary to uncover the underlying factors that govern these clustering patterns.

Prior studies have suggested that the bulk of K_V_2.1 channels heterologously expressed in Xenopus oocytes (*28*) or HEK293T cells (*19*) as well as endogenous K_V_2.1 in hippocampal neurons (*45*) and arterial myocytes (*15*) are in a nonconducting state. The prevailing view is that aggregation of K_V_2.1 channels into high density clusters is what renders most of these channels incapable of conducting K^+^ (45). Although our study does not address this issue comprehensively, at a minimum, our data suggest that K_V_2.1 conduction is not dependent on macro-clustering formation. Future studies should investigate whether the formation of K_V_2.1 micro-clusters may be sufficient to electrically silence these channels.

This is the first study to definitively demonstrate the structural role of K_V_2.1 clustering in regulating Ca_V_1.2 channel clustering and activity that occurs in channels of native cells. This is significant because the generally accepted view is that the functional impact of ion channel clustering is to exclusively concentrate ion conducting roles at specific sites. For example, Na^+^ channel clustering at nodes of Ranvier (*46*), neuronal Ca^2+^ channel clustering at active zones in presynaptic terminals (*47*), and skeletal muscle Ca^2+^ channels at SR Ca^2+^ release units (*48*). In the case of ventricular myocytes, it is concentrating voltage sensors at specific sites in the junctional dyad (*49, 50*). We propose that K_V_2.1 clustering is distinct in playing a role in modulating the localization and activity of an otherwise seemingly unrelated ion channel: Ca_V_1.2 channels. This functional impact of K_V_2.1 is due to the density-dependent cooperative gating that is an intrinsic property of Ca_V_1.2 channels (*51*).

Remarkably, the overall impact of K_V_2.1_S590A_ expression is that the differences between the I_Ca_ amplitude of wild-type male and female myocytes were eliminated in myocytes expressing the K_V_2.1S590A mutation, similar to what we observed in homozygous K_V_2.1 knockout mice (*15*). Thus, declustering K_V_2.1 channels appears to have the same impact as fully eliminating K_V_2.1 expression on Ca_V_1.2 clustering and activity in male and female myocytes. As our work also suggests that in arterial myocytes the conductive function of K_V_2.1 channels is independent of the degree of its clustering, in our model it is the extent of K_V_2.1 clustering that is the key determinant of the sex-specific differences in Ca^2+^ influx observed in these cells.

To conclude, we propose a model by which K_V_2.1 serves a structural role in promoting Ca_V_1.2 channel clustering and activity in a sex-dependent manner. Of note, K_V_2.1_S590A_ mutation reduced Ca_V_1.2 clustering and function in female myocytes but had no effect on male myocytes. K_V_2.1 clustering is not necessary for K_V_2.1 channel function however, K_V_2.1 macro-clusters alter Ca_V_1.2 channel organization. Together, our data suggest that the interactions between K_V_2.1 and Ca_V_1.2 are crucial for sex-based differences in arterial smooth muscle physiology.

## Materials and Methods

### Generation of the CRISPR/Cas9-edited K_V_2.1_S590A_ (KCNB1 S590A) knock-in mouse

The KCNB1 S590A mutation changes a AGC codon to GCC in Exon 2, thus converting a serine to an alanine (S590A) in the K_V_2.1 polypeptide. The knock-in mouse was generated in collaboration with the UC Davis Mouse Biology Program by using Crispr/CAS mediated homology directed repair. KCNB1 S590A mice were generated by introducing a mixture of gRNA (15 ng/L), single-stranded oligodeoxynucleotide (ssODN) repair template and Cas9 protein (30 ng/μL) by pronuclear microinjection into C57BL/6J mouse zygotes. Twenty zygotes were injected and implanted into the oviducts of one surrogate dam. A total of 6 pups were born, and genomic DNA was extracted from tail biopsies followed by PCR amplification using a specific primer set to identify a single male founder (F0). DNA-Seq analysis was used to confirm the mouse genotype. The correctly integrated single mutant F0 male mouse was further backcrossed with WT C57BL/6J female mice to produce offspring (F1) followed by intercrossing for two additional generations to obtain KCNB1 S590A heterozygotes which were used for breeding. Heterozygous and homozygous mutants were identified by a PCR genotyping protocol.

### Animals

Mice were euthanized with a single, lethal dose of sodium pentobarbital (250 mg/kg) intraperitoneally. All experiments were conducted in accordance with the University of California Institutional Animal Care and Use Committee guidelines.

### Arterial myocyte isolation

Third and fourth order mesenteric arteries were carefully cleaned of surrounding adipose and connective tissue, dissected out, and placed in ice-cold dissecting solution (Mg^2+^-PSS; 5 mM KCl, 140 mM NaCl, 2mM MgCl_2_, 10 mM glucose, and 10 mM HEPES) adjusted to pH 7.4 with NaOH. Arteries were first placed in dissecting solution supplemented with 1.23 mg/ml papain (Worthington Biochemical, Lakewood, NJ) and 1 mg/ml DTT at 37°C for 14 min. This was immediately followed by a five-min incubation in dissecting solution supplemented with 1.6 mg/ml collagenase H, 0.5 mg/ml elastase (Worthington Biochemical), and 1 mg/ml trypsin inhibitor from *Glycine max* at 37°C. Arteries were rinsed three times with dissection solution and single cells obtained by gentle trituration with a wide-bore glass pipette. Myocytes were maintained at 4°C until used.

### HEK293T cell culture and transfection

HEK293T (AATC #CRL-3216) cells were cultured in Dulbecco’s Modified Eagle Medium (Gibco #11955) supplemented with 10% fetal bovine serum (Gibco #26140) and 1% penicillin/streptomycin (Gibco #15140122) and maintained at 37°C in a humidified 5% CO_2_ atmosphere. Cells were transiently transfected using JetPEI (Polyplus Transfection #101000053) according to manufacturer’s protocol and passaged 24 hours later onto 25 mm square #1.5 coverslips or 18 mm square collagen coated #1.5 coverslips (Neuvitro Corporation #GG-18-15-Collagen) for GSD experiments. Plasmids encoding DsRed-Kv2.1_WT_, DsRed-Kv2.1_S586A_, DsRed-Kv2.1_P404W_, and DsRed-Kv2.1_P404W_, _S586A_ were previously described(*17, 21*). mScarlet-tagged versions of these plasmids were generated by GenScript, replacing the sequence encoding dsRed with sequence encoding mScarlet(*52*). For the bimolecular fluorescence experiments, cells were transfected with the pore-forming subunit of the rabbit Ca_V_1.2 (α1c, kindly provided by Dr. Diane Lipscombe; Brown University, Providence, RI) with the carboxy tail fused to either the N-fragment (VN) or the C-fragment (VC) of the Venus protein (27097, 22011; Addgene, Cambridge, MA), auxiliary subunits Ca_V_α_2_δ, Ca_V_β_3_ (kindly provided by Dr. Diane Lipscombe, Brown University, Providence, RI) and either DsRed-K_V_2.1_P404W_ or DsRed-K_V_2.1_P404W, S586A_-dsRed. HEK293T cells were transfected with Ca_V_1.2-VN, Ca_V_1.2-VC, Ca_V_α_2_δ, Ca _V_β_3_ and DsRed-K_V_2.1-dsRed in a 1.0:1.0:1.0:1.5:0.4 ratio.

### Live cell confocal imaging

HEK293T cells transfected with 200 ng of mScarlet-Kv2.1_WT_ or mScarlet-Kv2.1_S586A_ and seeded onto 25-mm square 1.5 coverslips approximately 16 hours before experiments. Imaging was performed in Tyrode III solution consisting of (in mM) 140 NaCl, 5.4 KCl, 1 MgCl_2_, 1.8 CaCl_2_, 5 HEPES, and 5.5 glucose, pH 7.4 with NaOH. Cells were imaged with an Olympus Fluoview 3000 confocal laser-scanning microscope equipped with an Olympus Plan-Apochromat 60x oil immersion lens (NA = 1.40). Image stacks were analyzed using Imaris software.

Stacks of images were analyzed using Imaris 10 (Andor, Belfast). Briefly, K_V_2.1-associated mScarlet signal was mapped to x/y/z centroid co-ordinates in each image stack using the Spots tool. Spots were assigned to all signal surpassing a fixed signal threshold and restricted to puncta greater than 100 nm (x/y) and 150 nm diameter (z), such that any bright signal with a volume greater than two voxels was identified as a K_V_2.1 cluster. ‘Region Growing’ was utilized (with a fixed manual threshold) to apply variable sizing to K_V_2.1 Spots, in line with the volume and brightness of mScarlet puncta. Finally, the Cell segmentation function was used to estimate cell boundaries based on low-intensity mScarlet signal and obtain an approximate cell volume.

### K_V_2.1 immunofluorescence immunocytochemistry

Immunofluorescence labeling was performed on freshly dissociated arterial myocytes. Cells were left to adhere for one hour at room temperature prior to fixation, fixed with 4% formaldehyde (Electron Microscopy Sciences #50980487) diluted in phosphate-buffered saline (PBS) (Fisher Scientific, Hampton, NH) for 15 minutes at room temperature, washed, and incubated with 50 mM glycine (BioRad, Hercules, CA) for 10 min to reduce aldehydes. The surface membrane was stained with wheat germ agglutinin (WGA) Alexa Fluor 488 (1 μM, ThermoFisher #W11261) for 10 minutes at room temperature followed by washing. Cells were then incubated in blocking buffer made of 3% w/v bovine serum albumin and 0.25% Triton X-100 in PBS, followed by incubation with mouse anti-Kv2.1 (mAb K89/34; RRID: AB_2877280; NeuroMab, Davis, CA, 1:200) diluted in blocking buffer for one hour at room temperature or overnight at 4°C. Myocytes were washed, incubated at room temperature for one hour with Alexa Fluor 647-conjugated donkey anti-mouse IgG diluted in blocking buffer (2 μg/ml, Molecular Probes, Cat #A31571) followed by washes in PBS. For experiments investigating K_V_2.1 phosphorylation state, double labeling was performed with the mouse anti-K_V_2.1 pS590 phosphospecific mAb L100/1(*30*) together with rabbit anti-Kv2.1 (KC(*23*); Trimmer laboratory, RRID:AB_2315767; 1:100). Myocytes were washed, incubated at room temperature for one hour with Alexa Fluor 568-conjugated goat anti-mouse IgG (2 μg/ml, Molecular Probes, Cat #A11004) and Alexa Fluor 647-conjugated donkey anti-rabbit IgG (2 μg/ml, Molecular Probes, Cat #A31571) diluted in blocking buffer followed by washes in PBS. All washes were performed with PBS three times for 10 minutes. Coverslips were mounted onto microscope slides in Vectashield mounting medium (Vector Labs) and sealed with clear nail polish. Images were collected on a Dragonfly 200 spinning disk confocal (Andor), coupled to a DMi* Leica microscope (Leica, Wetzlar, Germany) equipped with a 60x oil immersion objective (NA = 1.40) and acquired using an Andor iXon EMCCD camera. Images were collected via Fusion software, in optical planes with a z-axis of 0.13 μm/step. Image files were analyzed using Imaris.

Image stacks were segmented and analyzed in Imaris 10. WGA-488 signal was background-subtracted and a fixed threshold applied to consistently map the plasma membrane, using the Surfaces tool. Alexa Fluor-647 signal (denoting K_V_2.1 puncta) was assessed using the Spots tool, as described above. Spots marking K_V_2.1 clusters were categorized into internal and plasma membrane-restricted components, with the latter utilized for analysis.

### Super-resolution microscopy

HEK293T cells transfected with 200 ng mScarlet-Kv2.1_WT_ or mScarlet-Kv2.1_S586A_-mScarlet and arterial myocytes were plated onto collagen coated glass coverslips (Neuvitro Corporation, #GG-18-1.5-Collagen) followed by fixation with 3% formaldehyde and 0.1% glutaraldehyde diluted in PBS for 15 min at room temperature. After washing with PBS, cells were incubated with 50 mM glycine (BioRad, Hercules, CA) for 10 min to quench aldehydes. Cells were washed and incubated for one hour at room temperature with a blocking buffer made with 3% w/v BSA and 0.25% Triton X-100 in PBS. Cells were then incubated with either mouse anti-Kv2.1 (HEK293T experiments, mAb K89/34; RRID: AB_2877280; UC Davis/NIH Neuromab Facility, Davis, CA; 1:20) or mouse anti-Ca_V_1.2 (arterial myocytes experiments, mAb L57/23; RRID: AB_2802123; 1:5). After extensive washings with PBS (three quick washes followed by three 30-min washes), cells were incubated at room temperature for one hour with Alexa Fluor 647-conjugated goat anti-mouse diluted in blocking buffer to a concentration of 2 μg/ml and afterwards extensively washed with PBS.

The imaging buffer contained 10 mM MEA, 0.56 mg/ml glucose oxidase, 34 μg/ml catalase, and 10% w/v glucose in TN buffer (200 mM Tris-HCl pH 8, 10 mM NaCl). A super resolution ground state deletion system (SR-GSD, Leica, Wetzlar, Germany) based on stochastic single-molecule localization was used to generate super-resolution images of Ca_V_1.2 and K_V_2.1 labeling. The Leica SR-GSD is a Leica DMI6000B TIRF microscope system equipped with a 160x HCX Plan-Apochromat (NA 1.43) oil-immersion lens and an EMCCD camera (iXon3 897, Andor Technology, Belfast, United Kingdom). Fluorophores were excited with a 642 nm laser (used for both pumping to the dark state and image acquisition). For all experiments, the camera was running in frame-transfer mode at a frame rate of 100 Hz (11 ms exposure time). Fluorescence was detected through Leica high power TIRF filter cubes (488 HP-T, 532 HP-T, 642 HP-T) with emission band-pass filters of 505-605 nm, 550-650 nm, and 660-760 nm. A total of 35,000 images were collected per cell and used to construct the super resolution localization images. Fluorescence signals in each image were fit with a 2D Gaussian function which localized the coordinates of centroids of single molecule fluorescence within the LASAF software (Leica). Images were rendered at 20 nm/pixel (normalized Gaussian mode), threshold (# photons/event) using the GSD software and exported as binary TIF images. Particle analyses were determined in ImageJ. Representative images were rendered down to 2 nm for visualization purposes.

To accomplish the Gaussian blur, the GSD generated pixel in an image was replaced by a weighted average of 200 nm of its neighboring pixels. The amount of blur applied to the image was controlled by the size of the kernel, which determines the radius of the neighboring pixels used in the calculation, such that the larger the kernel, the more pixels are included in the calculation, and the stronger the blur effect.

### Quantitative PCR

Total RNA was isolated using the RNeasy Mini Kit (Qiagen) as per manufacturer’s instructions. Isolated mRNA was then reverse transcribed using the AffinityScript qPCR cDNA Synthesis Kit (Agilent) following manufacturer’s protocol. Quantitative PCR (qPCR) analysis was performed using a QuantStudio 7 Pro Real-time PCR System (Applied Biosystems), using PowerUP SYBR Green Master Mix (Thermo Fisher Scientific) as the fluorescence probe. The cycling conditions were 50°C for 2 minutes and 95°C for 10 minutes, followed by 40 cycles of 95°C for 15 s and 56°C for 1 minute. A dissociation curve protocol (ramping temperatures between 60°C and 95°C) was added at the end to verify amplification specificity of each qPCR reaction.

Specific primers were designed in this experiment, including β-actin (NM_007393.5): sense nt (895-914): CCAGCCTTCCTTCTTGGGTA, antisense nt (989-967): AGAGGTCTTTACGGATGTCAACG; and Ca_V_1.2 (NM_009781.4): sense nt (5-23): CTGAAAGCAGAAGCTCGGA, antisense nt (181-163): CATTGTGGCTTCCAGTTGG. Primer efficiencies were tested to be in between 90% and 110%. The relative abundance of Ca_V_1.2 transcript was normalized to β-actin transcript expression.

### Proximity Ligation Assay

A Duolink In Situ PLA kit (Sigma) was used to detected K_V_2.1-K_V_2.1 and K_V_2.1-Ca_V_1.2 complexes in freshly isolated mesenteric arterial myocytes. All protocols post incubation of primary antibodies were followed in accordance with the manufacturer’s instructions. Briefly, cells were plated on glass coverslips and allowed to adhere for 1 hour at room temperature. Cells were fixed with 4% paraformaldehyde for 20 minutes, quenched in 10mM glycine for 15 min, washed in PBS two times for three minutes, and permeabilized 20 minutes in 0.1% Triton X-100. After blocking for 1 hour at 37°C in Duolink Blocking Solution, cells were incubated overnight at 4°C using the following primary antibodies: mouse anti-Kv2.1 (mAb K89/34; RRID: AB_2877280; UC Davis/NIH Neuromab Facility, Davis, CA; 1:200), rabbit anti-Kv2.1 (KC(*23*); RRID:AB_2315767; 1:100) and rabbit anti-Ca_V_1.2. The anti-Ca_V_1.2 rabbit polyclonal antibody “Cav1.2 II-III” was generated by immunizing two New Zealand white rabbits with a His-tagged recombinant protein fragment corresponding to a.a. 785-900 of mouse Ca_V_1.2 (accession number Q01815). Antibodies were affinity purified from serum on nitrocellulose strips containing the Cav1.2 II-III His-tagged recombinant protein fragment following the method of Olmsted(*53*). Cells incubated with only one primary antibody served as negative controls. Secondary oligonucleotide-conjugated antibodies (PLA probes: anti-mouse MINUS and anti-rabbit PLUS) were used to detect K_V_2.1 and Ca_V_1.2 interactions. Fluorescent signal was detected using an Olympus FV3000 confocal microscope equipped with a 60x oil immersion lens (NA = 1.40). Images were collected with a z-axis of 0.5 μm/step optical planes. Stacks of images were combined in ImageJ and used for analysis of puncta/μm^2^ per cell.

### Patch-clamp electrophysiology

All electrophysiological recordings were acquired at room temperature using an Axopatch 200B amplifier and Digidata 1440 digitizer (Molecular Devices, Sunnyvale, CA). Borosilicate patch pipettes were pulled and polished to resistances of 3-6 MΩ using a micropipette puller (model P-97, Sutter Instruments, Novato, CA).

I_Kv2.1_ was measured in arterial myocytes using conventional whole-cell voltage-clamp electrophysiology at a frequency of 50 kHz and low-pass filter of 2 kHz. Cells were continuously perfused with an external solution containing (in mM) 130 NaCl, 5 KCl, 3 MgCl_2_, 10 glucose, and 10 HEPES adjusted to pH 7.4 with NaOH. Micropipettes were filled with an internal solution containing (in mM) 87 K-aspartate, 20 KCl, 1 CaCl_2_, 1 MgCl_2_, 5 MgATP, 10 EGTA, and 10 HEPES pH 7.2 with KOH. A liquid junction potential of 13 mV was corrected for offline. To measure current-voltage relationships, cells were subjected to a series of 500 ms test pulses increasing from -70 mV to +70 mV. In order to isolate the RY785-sensitive Kv2 current, cells were first bathed and recorded in external solution. Cells were then exposed to 1μM RY785 (MedChemExpress) to inhibit Kv2.1 activity. RY785-sensitive currents were calculated by subtracted the RY785 exposed traces from the composite I_K_ traces.

I_Ca_ was measured in isolated arterial myocytes using conventional whole-cell electrophysiology. Currents were measured at a frequency of 50 kHz and low-pass filtered at 2 kHz. Myocytes were continuously bathed in an external solution with (in mM) 115 NaCl, 10 TEA-Cl, 0.5 MgCl_2_, 5.5 glucose, 5 CsCl, 20 BaCl_2_, and 10 HEPES adjusted to a pH of 7.4 using CsOH. Micropipettes were filled with (in mM) 20 CsCl, 87 aspartic acid, 1 MgCl_2_, 10 HEPES, 5 MgATP, and 10 EGTA adjusted to pH 7.2 via CsOH. A voltage error of 9.4 attributed to the liquid junction potential of the recording solutions was corrected for offline. Cells were exposed to a series of 300 ms depolarizing pulses from a holding potential of -70 mV to test potentials ranging from -70 mV to +60 mV to attain current-voltage relationships.

### Bimolecular fluorescence complementation

Spontaneous interactions of Ca_V_1.2 channels were assayed using biomolecular fluorescence complementation. HEK293T cells were transfected with Ca_V_1.2 channels tagged at their C-terminus to either a non-fluorescent N-(VN(1-154, I152L)) or C-terminal (VC(155-238, A206K)) halves of a ‘split’ Venus fluorescent protein. When Ca_V_1.2-VN and Ca_V_1.2-VC are brought close enough together to interact, the full Venus protein can fold into its functional, fluorescent conformation. The scale of Venus fluorescence emission can therefore be an indicator of Ca_V_1.2-Ca_V_1.2 interactions. Venus fluorescence was monitored using TIRF microscopy

For whole-cell current recordings from HEK293T cells, pipettes were filled with a solution containing (mM) 84 Cs-aspartate, 20 CsCl, 1 MgCl_2_, 10 HEPES, 1 EGTA, and 5 MgATP adjusted to pH 7.2 using CsOH. HEK293T cells were continuously perfused with an external solution comprising of (in mM) 5 CsCl, 10 HEPES, 10 glucose, 140 NMDG, 1 MgCl_2_, and 20 CaCl_2_ with a pH of 7.4 (HCl).

### Ca_V_1.2 sparklets

Ca^2+^ sparklets were recorded using a through-the-lens TIRF microscope built around an inverted microscope (IX-70; Olympus) equipped with a Plan-Apochromat (60X; NA 1.49) objective (Olympus) and an electron-multiplying charge-coupled device (EMCCD) camera (iXON; Andor Technology, UK). Myocytes were loaded via the patch pipette with a solution containing (in mM) 0.2 Fluo-5F (Invitrogen # F14221), 87 Cs-aspartate, 20 CsCl, 1 MgCl_2_, 5 MgATP, 10 HEPES, 10 EGTA, pH 7.2 with CsOH. Cells were perfused in an external solution containing 140 NMDG, 5 CsCl, 51 MgCl_2_, 10 glucose, 10 HEPES, 2 CaCl_2_, pH 7.4 with HCl. After obtaining a GΩ seal in 2mM Ca^2+^ external solution, the cell was broken into and allowed to dialyze for 3 minutes. The external solution was exchanged with a solution containing (in mM) 120 NMDG, 5 CsCl, 1 MgCl_2_, 10 glucose, 10 HEPES, 20 CaCl_2_, pH 7.4 with HCl. Images for the detection of sparklets were recorded at a frequency of 100 Hz using TILL Image software. Cells were held to a membrane potential of −70 mV using the whole-cell configuration of the patch-clamp technique. Sparklets were automatically detected and later analyzed using custom software (Source code 2) written in MATLAB (RRID:<u>SCR_001622</u>) as previously described (*10*).

### In silico modeling

Simulations were performed using the Sato et al. (*9*) and Hernandez-Hernandez et al. (*31*) mathematical models for cluster formation and smooth muscle electrophysiology, respectively.

### Chemicals and statistics

All chemical reagents were acquired from Sigma-Aldrich (St. Louis, MO) unless otherwise stated. Data was expressed as mean ± SEM and analyzed using GraphPad Prism software. Statistical significance was determined using appropriate paired or unpaired T-tests, non-parametric tests, or one-way analysis of variance (ANOVA). P < 0.05 was considered statistically significant and denoted by * in the figures.

## Acknowledgments

We thank Dellaney Rudolph-Gandy and Ernesto Javier Rivera for technical assistance.

## Funding

This work was supported by grants from:

US National Institutes of Health HL085686

US National Institutes of Health HL128537

US National Institutes of Health HL144071

US National Institutes of Health NS114210

US National Institutes of Health 1OT2OD026580

Amazon AWS Cloud Credits for Research.

## Author contributions

Conceptualization: CM, LFS, JST

Methodology: CM, LFS, NCV, JST

Investigation: CM, SCO, DM, GHH, PR, ZF, DS

Formal Analysis: CM, SCO, DM, PR, ZF, LFS

Visualization: CM

Data Curation: GHH, DS

Software: GHH, DS

Supervision: LFS, CEC, JST

Writing—original draft: CM, LFS

Writing—review & editing: CM, SCO, DM, GHH, PR, ZF, NCV, JST, LFS

## Data and materials availability

All data are available in the main text or the supplementary materials.

