## Supplemental Figures for "The formation of K_V_2.1 macro-clusters is required for sex-specific differences in L-type Ca_V_1.2 clustering and function in arterial myocytes"

Collin Matsumoto *et al.*

**This file includes:**

Figs. S1 to S4

**Fig. S1.**
**The distributions of $K_V2.1_{WT}$ and $K_V2.1_{S586A}$ in HEK293T cells could be explained by a stochastic self-assembly mechanism.**

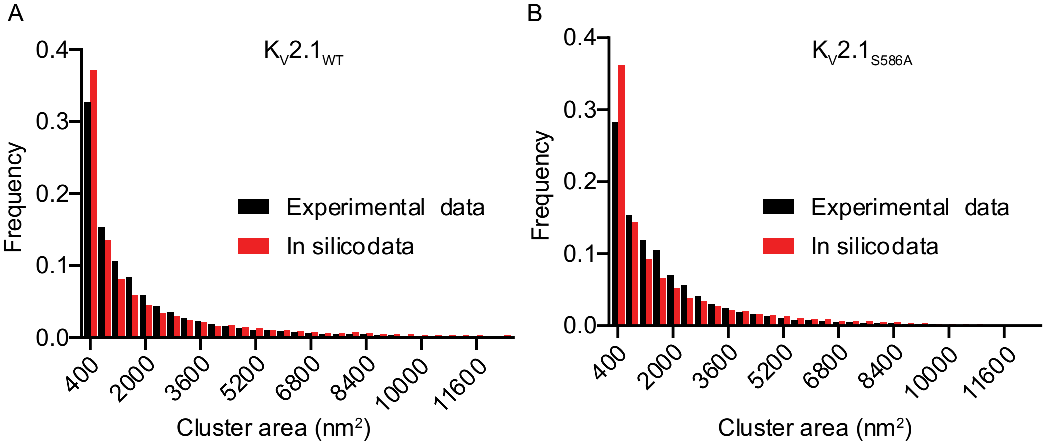


C

| Channel | $P_n$ | $P_g$ | $P_R$ | Mean experimental cluster size (nm$^2$) | Mean *in silico* cluster size (nm$^2$) | Mean experimental cluster density (clusters/µm$^2$) | Mean *in silico* cluster density (clusters/µm$^2$) |
|---|---|---|---|---|---|---|---|
| $K_V2.1_{WT}$ | 0.00050303 | 0.16532 | 0.001 | 2634 ± 143 | 2634.1 | 41 ± 4.7 | 40.389 |
| $K_V2.1_{S586A}$ | 0.000098767 | 0.050355 | 0.001 | 1860 ±104 | 1867.3 | 29 ± 3.7 | 28.758 |

(**A**) Histograms of experimental (black bars) and simulated (red bars) cluster area distributions as a relative frequency of $K_V2.1_{WT}$ in HEK293T cells. (**B**) Histograms of experimental (black bars) and simulated (red bars) cluster area distributions as a relative frequency of $K_V2.1_{S586A}$ in HEK293T cells. (**C**) Summary of experimental and *in silico* data.

**Fig. S2.**

**Supplemental Figure 2. K$^+$ currents in K$_V$2.1$_{WT}$, K$_V$2.1$^{-/-}$ null and K$_V$2.1$_{S590A}$ myocytes.**

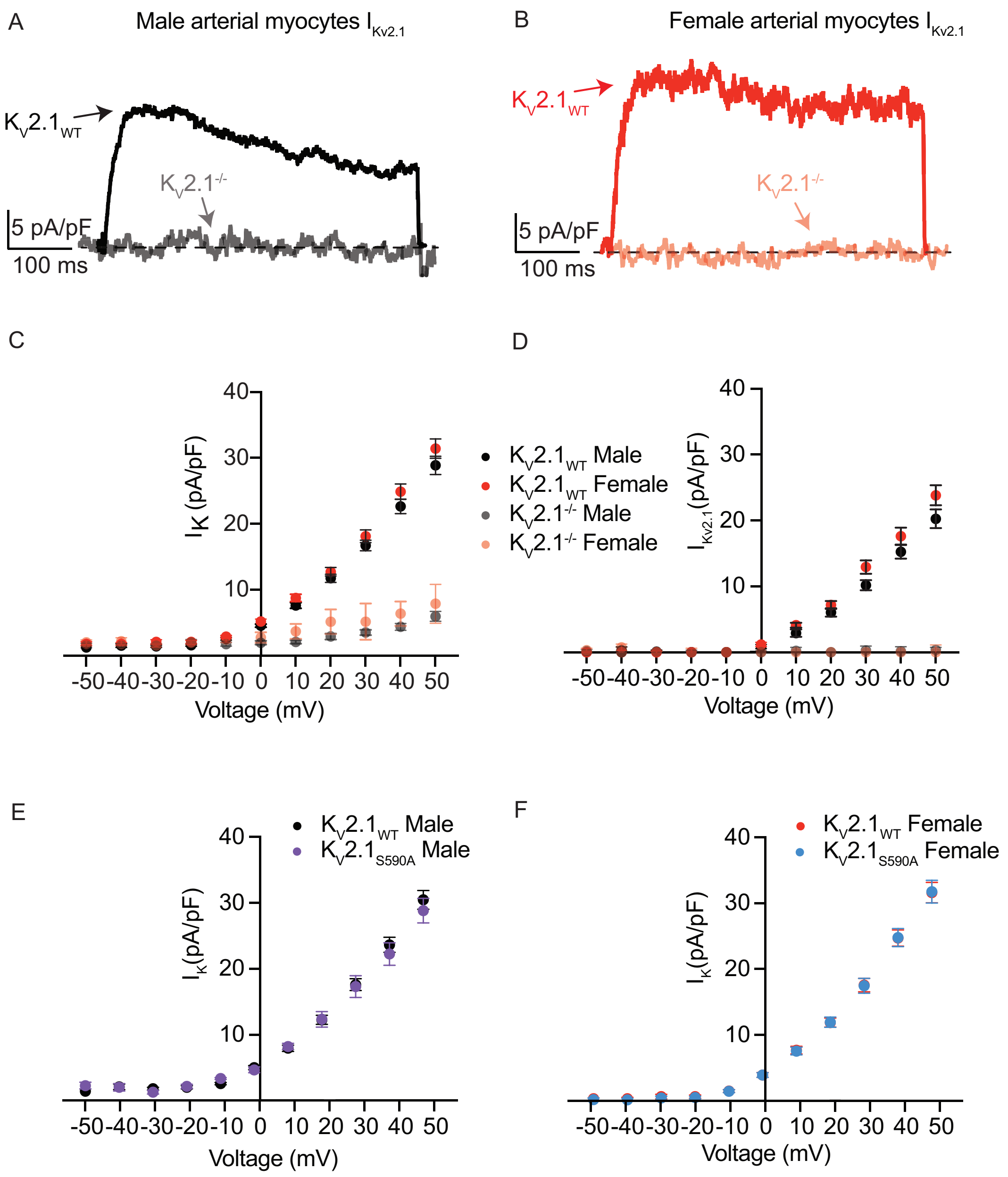


(**A**) Representative RY785-sensitive traces at +50 mV from K$_V$2.1$_{WT}$ (black) and K$_V$2.1$^{-/-}$ null (gray) males myocytes. (**B**) Representative traces at +50 mV from K$_V$2.1$_{WT}$ (red) and K$_V$2.1$^{-/-}$ null (pink) female myocytes. (**C**) IV relationship of total K$^+$ current (I$_K$) recorded from K$_V$2.1$_{WT}$ male (black), K$_V$2.1$^{-/-}$ null male (gray), K$_V$2.1$_{WT}$ female (red), and K$_V$2.1$^{-/-}$ null female (pink) myocytes. (**D**) IV relationship of RY785-sensitive (Kv2.1) currents recorded from K$_V$2.1$_{WT}$ male (black), K$_V$2.1$^{-/-}$ null male (gray), K$_V$2.1$_{WT}$ female (red), and K$_V$2.1$^{-/-}$ null female (pink) myocytes. (**E**) IV relationship of total K$^+$ current (I$_K$) recorded from K$_V$2.1$_{WT}$ male (black) and K$_V$2.1$_{S590A}$ male (purple) myocytes. (**F**) IV relationship of total K$^+$ current (I$_K$) recorded from K$_V$2.1$_{WT}$ female (red) and K$_V$2.1$_{S590A}$ female (blue) myocytes.

**Fig. S3.**
**Supplemental Figure 3. $K_V2.1$ and $Ca_V1.2$ are decreased in female $K_V2.1_{S590A}$ myocytes.**

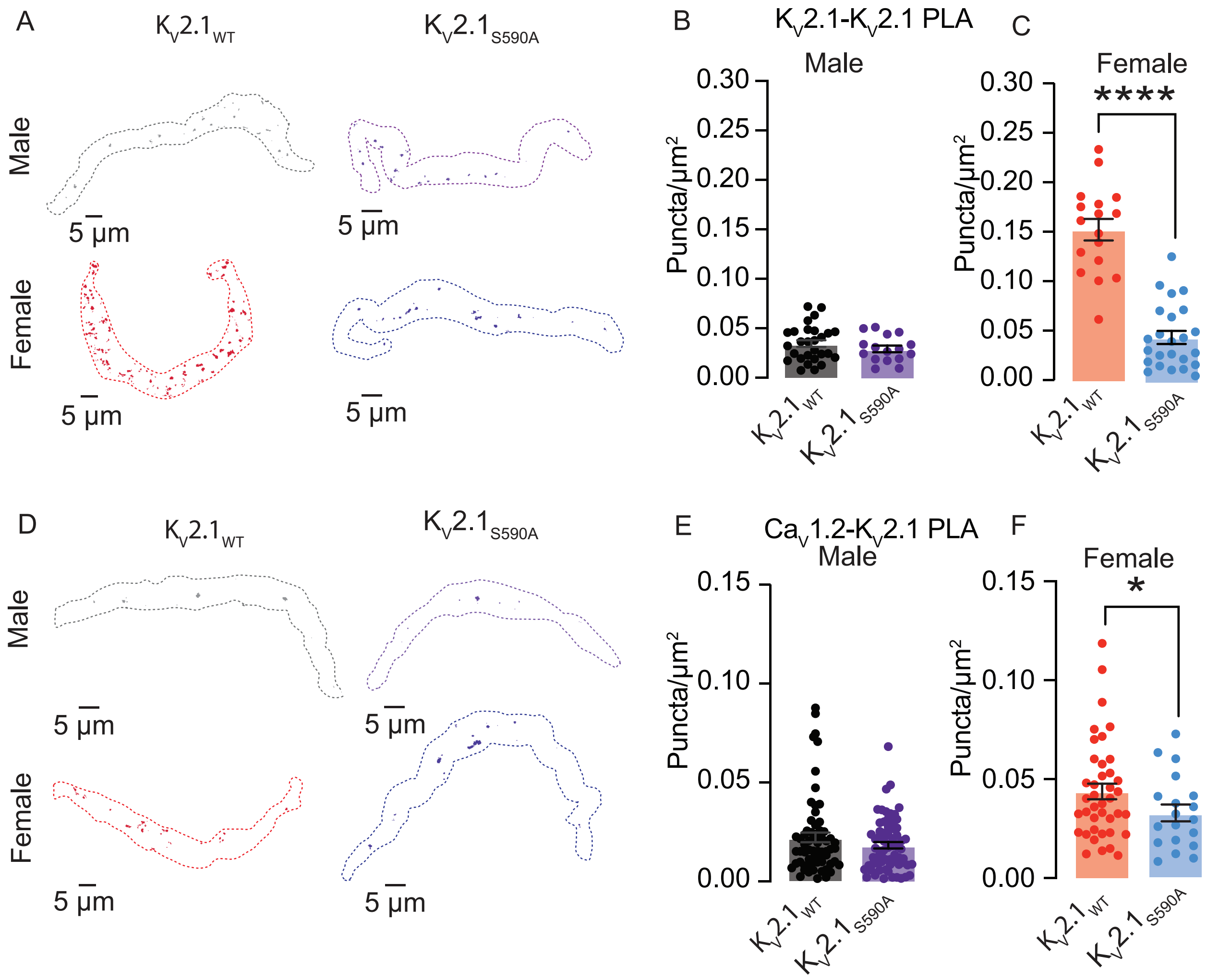


(**A**) PLA images of representative $K_V2.1$-$K_V2.1$ channel interactions in arterial myocytes. Quantification of $K_V2.1$-$K_V2.1$ PLA puncta/µm$^2$ in male (**B**) and female (**C**) myocytes. (**D**) PLA images of representative $K_V2.1$-$Ca_V1.2$ channel interactions in arterial myocytes. Quantification of $K_V2.1$-$Ca_V1.2$ PLA puncta/µm$^2$ in male (**E**) and female (**F**). *P<0.05, **P<0.01, ***P<0.001, ****P<0.0001. Error bars indicate mean ± SEM.

**Fig. S4**

**Supplemental Figure 4 The distributions of $K_V2.1_{WT}$ and $K_V2.1_{S590A}$ in arterial myocytes could be explained by a stochastic self-assembly mechanism.**

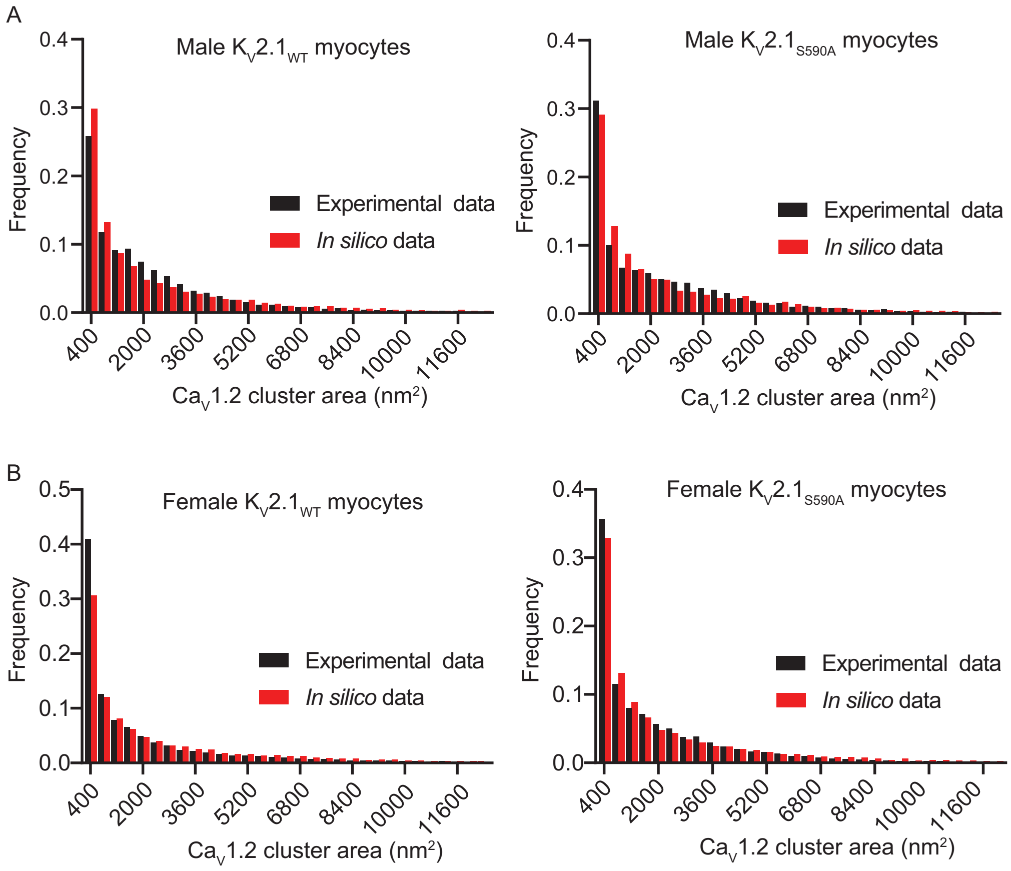


C

| Arterial Myocyte CaV1.2 | $P_n$ | $P_g$ | $P_R$ | Mean experimental cluster size ($nm^2$) | Mean *in silico* cluster size ($nm^2$) | Mean experimental cluster density (clusters/$\mu m^2$) | Mean *in silico* cluster density (clusters/$\mu m^2$) |
|---|---|---|---|---|---|---|---|
| WT male | 0.000056159 | 0.083203 | 0.001 | 2249 ± 54.53 | 2567.3 | 14 ± 1.2 | 14.062 |
| $K_V2.1_{S590A}$ male | 0.000047767 | 0.086086 | 0.001 | 2345 ± 81.84 | 2574.6 | 12 ± 2.2 | 12.176 |
| WT female | 0.000047826 | 0.10793 | 0.001 | 3098 ±163.8 | 2989.9 | 11 ± 1.1 | 10.881 |
| $K_V2.1_{S590A}$ female | 0.000043453 | 0.067511 | 0.001 | 2175 ± 91.72 | 2327.7 | 13 ± 1.8 | 13.32 |

(**A**) Histograms of experimental (black bars) and simulated (red bars) $Ca_V1.2$ cluster area distributions as a relative frequency of $K_V2.1_{WT}$ (left) and $K_V2.1_{S590A}$ (right) in male arterial myocytes. (**B**) Histograms of experimental (black bars) and simulated (red bars) $Ca_V1.2$ cluster area distributions as a relative frequency of $K_V2.1_{WT}$ (left) and $K_V2.1_{S590A}$ (right) in female arterial myocytes. (**C**) Summary of experimental and *in silico* data.
